# Scalable recorders of cellular calcium dynamics and molecular interactions across time

**DOI:** 10.64898/2026.11.13.765775

**Authors:** Yixiao Yan, Changyang Linghu

## Abstract

Cellular processes are mediated by dynamic biochemical events in their native context, including changes in intracellular ion concentrations and transient interactions between molecules. Studying these processes across large cell populations requires recording their dynamics across time in single cells, simultaneously and at scale. Here, we introduce split-1POK, a split protein tape monomer whose two halves reconstitute when the domains fused to them interact. The reconstituted monomer is incorporated into the protein tape as it elongates across time in live cells, enabling scalable, spatiotemporally resolved endpoint readout under conventional light microscopy. Split-1POK provides a programmable scaffold for new recorders in the CytoSense family^1^, our previously introduced framework in which the same cytosol-localized monomer senses cellular activity and writes it along protein tape recorders. Unlike transcription-based activity recorders, which encode the signal through new gene expression, CytoSense recorders act on monomers post-synthesis, so the activity signal is sensed and written directly in the cytosol without passing through transcription- or translation-associated cascades. Fusing calmodulin and its binding peptide to split-1POK, we built TCaMP (Tape monomer-CalModulin-Peptide), a genetically encoded calcium recorder (GECR) that measures calcium dynamics on a single, analog intensity scale along a continuous time axis, providing quantitative comparison of calcium levels over time within the same cell. In cultured human cells, TCaMP records transient calcium dynamics from transmembrane entry, receptor activation, and intracellular store release, detecting stimulations as brief as ten minutes. With hour-scale kinetics over days, TCaMP complements fluorescent calcium indicators by trading sub-second temporal resolution for scalable, long-term recording. Fusing conditional dimerization pairs to split-1POK yields recorders of molecular interaction histories in response to external stimuli. Split-1POK thus brings scalable, spatiotemporally resolved recording to intracellular calcium dynamics and molecular interactions across time.

## 1 Introduction

Calcium is a highly dynamic, universal second messenger regulating many essential cellular processes^2^. Decades of development in calcium indicators and calcium integrators have transformed the study of these processes in single cells and across cell populations, with trade-offs among spatial resolution, spatial scale, and temporal continuity. Real-time imaging of fluorescent calcium indicators, including synthetic dyes and genetically encoded indicators such as GCaMPs^3–5^, provides temporal resolution down to the millisecond scale, but their readout requires sustained optical access. That access is bounded in space by field-of-view size and light scattering and absorption in tissue, which limit the imaging volume at single-cell resolution, and in time by photobleaching, which limits recording duration. Genetically encoded calcium integrators, such as the CaMPARI and FLARE families^6,7^, enable retrospective, scalable readout of calcium activity integrated within a single defined time window, typically yielding one intensity value per cell. More recently, Caprola provides a scalable readout of calcium activity across discrete time windows, each defined by the supply of a spectrally distinct fluorescent HaloTag ligand and read out in its own spectral channel, such that activity in each window is reported on a separate intensity scale^8^. A “genetically encoded calcium recorder” (GECR) that records calcium dynamics on a single, analog intensity scale across a continuous time axis with both spatial scalability and single-cell precision has yet to be achieved.

Here, we introduce split-1POK, a programmable molecular scaffold for building intracellular activity recorders, and TCaMP, a genetically encoded recorder of calcium dynamics built on it (**Figure 1**). TCaMP continuously writes calcium dynamics on a single, analog intensity scale along elongating protein tape assemblies^9,10^ in live cells, for scalable endpoint in situ readout from single cells across a real-world time axis. The TCaMP tape monomer itself both senses calcium and becomes part of the tape record, acting in the cytosol throughout. In human U2OS cells, TCaMP records calcium dynamics with hour-scale kinetics over days, detecting calcium elevations from transmembrane entry, from G protein-coupled receptor (GPCR) activation, and from intracellular store release. TCaMP is sensitive to stimulations as brief as ten minutes and distinguishes the amplitude and timing of successive calcium events along the time axis. We also reprogrammed split-1POK to record molecular interaction histories of conditional dimerization pairs in response to external stimuli in live cells. Together, split-1POK and the recorders it enables establish a programmable framework for scalable intracellular recording of calcium dynamics, molecular interaction histories, and beyond.

**Figure 1.**
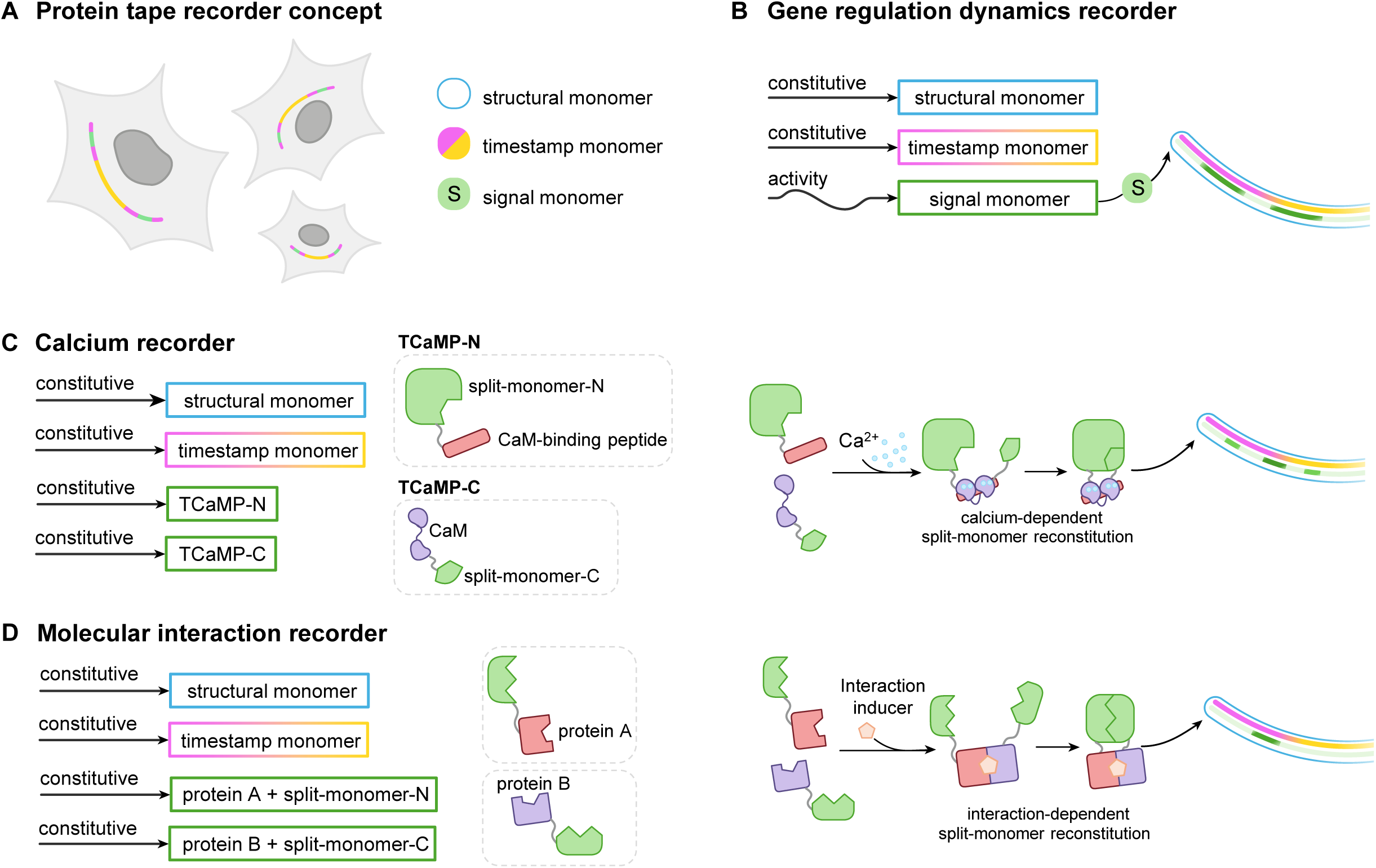
A modular split-monomer strategy enables protein tape recording of calcium dynamics and molecular interactions. (**A**) Concept of the protein tape recorder. An intracellular protein assembly encodes temporal cellular activity along a spatial axis with three genetically encoded components: structural monomers, timestamp monomers, and signal monomers, which differ by a short epitope tag. Recording is scalable across large cell populations, with tape records in single cells read out post hoc and in situ. (**B**) Existing protein tape recorders track gene regulation dynamics via transcriptionally regulated signal monomer expression, followed by subsequent signal monomer incorporation over time along a growing protein assembly established by constitutively expressed structural monomers. (**C** and **D**) The tape monomer was split into N- and C-terminal fragments that cannot independently incorporate into the growing protein tape. Cytosolic activity promotes the interaction between activity-sensing domains fused to the two fragments, which in turn modulates the reconstitution of the fragments into a full signal monomer that is subsequently incorporated into the tape. (**C**) A genetically encoded calcium recorder (TCaMP). The calcium-responsive interaction domains calmodulin (CaM) and a CaM-binding peptide were fused to the C- and N-terminal fragments of the split monomer, respectively. Intracellular Ca^2+^ promotes the interaction between CaM and the CaM-binding peptide, and thus the split-monomer reconstitution and subsequent incorporation into the growing protein tape. (**D**) Molecular interaction recorders. A pair of proteins (protein A and protein B) were fused to the N- and C-terminal fragments of the split monomer, respectively. The interaction between protein A and protein B, such as those inducible by a third molecule (interaction inducer), promotes split-monomer reconstitution and its subsequent incorporation into the growing protein tape.

## 2 Results

### 2.1 Development of split-1POK and recorders of molecular interactions

We first set out to couple protein tape recorders to molecular interactions. We reasoned that splitting the tape monomer into two separate domains and fusing each to one of a protein pair of interest would enable interaction-dependent tape incorporation, thereby preserving a temporal record of the corresponding signaling events. (**Figure 2A**). Such a recording mechanism would require that the two split-monomer domains reconstitute and incorporate onto the tape only when the two proteins of interest come into interaction.

**Figure 2.**
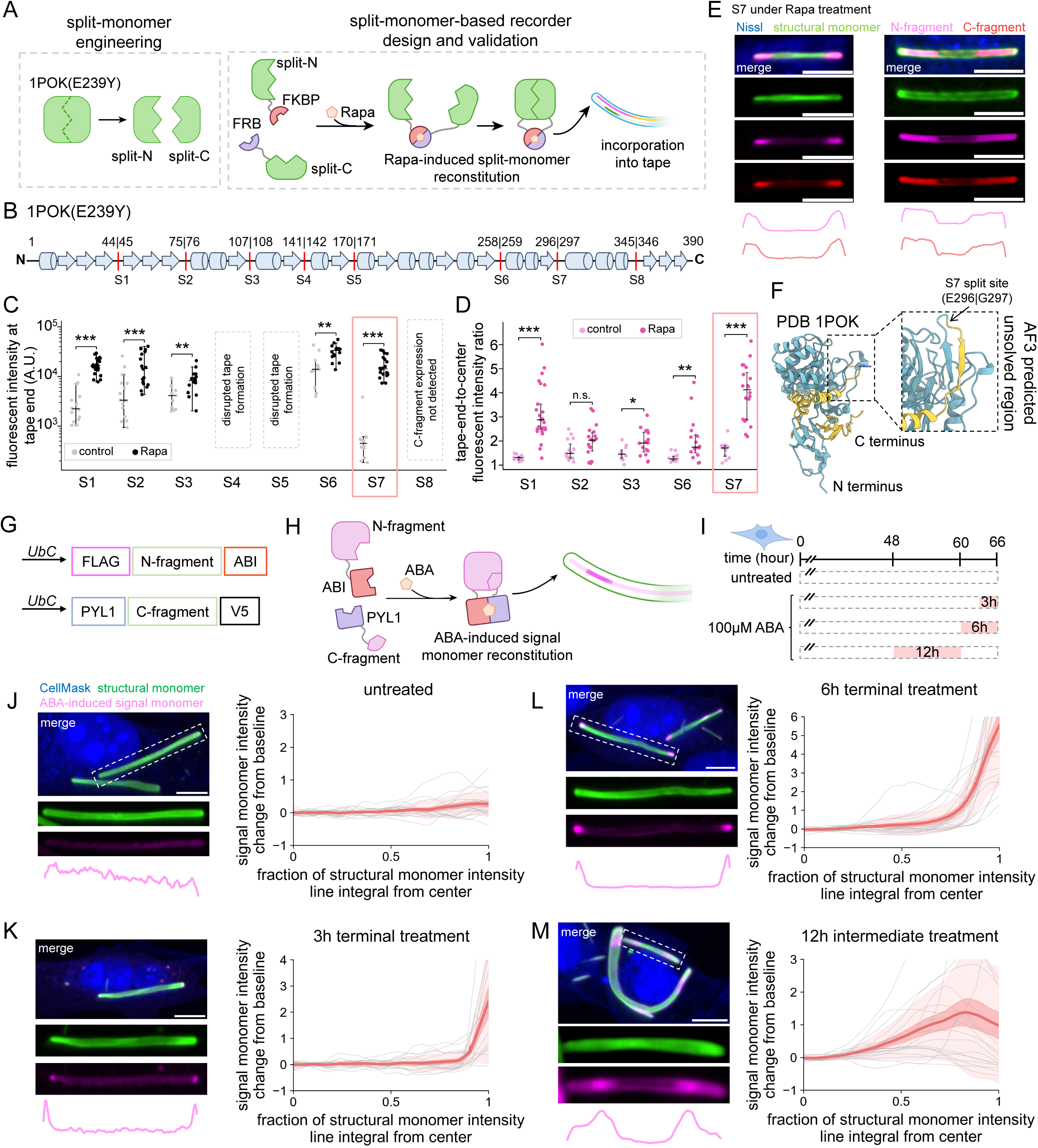
Split-1POK enables protein tape recording of molecular interactions. (**A**) Schematic of split-monomer engineering (left) and split-monomer-based recorder design and validation (right). The structural monomer of the protein tape, 1POK(E239Y), was split into N- and C-terminal fragments, which cannot independently incorporate into the tape. To validate the splitting designs, the rapamycin-inducible dimerization domains FKBP and FRB were fused to the N- and C-terminal fragments, respectively. Upon rapamycin (Rapa) treatment, FKBP–FRB dimerization promotes the reconstitution of the split monomer, enabling its subsequent incorporation into the growing tape. (**B**) Schematic representation of the secondary structure of 1POK(E239Y) aligned to the crystal structure of PDB: 1POK^26^. Cylinders represent α-helices, arrows represent β-strands, and connecting lines represent loop regions. Red vertical lines indicate the eight engineered split sites (S1–S8), with their corresponding amino acid positions labeled above. (**C**) Quantification of raw fluorescence intensity at tape ends in U2OS cells co-transfected with CytoTape-vivo and rapamycin-inducible split-monomer constructs, with (black) or without (gray) rapamycin (Rapa) treatment. Rapamycin was administered at 48 h for 2 hours after transfection before fixation at 72 h. Fluorescence intensity was measured using the epitope tag fused to the N-terminal fragment of each split monomer. Dots represent individual tapes, horizontal lines indicate medians, and whiskers extend to the most extreme values within 1.5 × IQR of the first and third quartiles; plotted in log scale. The tapes (cells) number in the control/Rapa groups are *n* = 14/25, 15/18, 11/16, 12/16, 12/20 for S1, S2, S3, S6, S7, respectively. Throughout this figure: the selected split site, S7, is highlighted in red. n.s., not significant; *, P < 0.05; **, P < 0.01; ***, P < 0.001; two-sided Mann-Whitney U test, with Holm correction for multiple comparisons. (**D**) Quantification of tape end-to-center fluorescence intensity ratio for five split-monomer designs exhibiting detectable expression, with (darker magenta) or without (lighter magenta) rapamycin (Rapa) treatment (median, IQR), measured using the epitope tag fused to the N-terminal fragment. The tapes (cells) number in the control/Rapa groups are *n* = 14/25, 15/18, 11/16, 12/16, 12/20 for S1, S2, S3, S6, S7, respectively. (**E**) Representative confocal images and the corresponding fluorescent intensity profiles for N- and C-fragments for the selected split site, S7. Scale bars, 10 μm. (**F**) Structural representation of 1POK(E239Y) aligned to the crystal structure of PDB: 1POK, with the N- and C-terminal regions colored in blue and yellow, respectively. The enlarged view highlights the selected split site S7 (E296|G297), located within a structurally unresolved region of the crystal structure, with the missing segment modeled using AlphaFold 3 (AF3)^27^. (**G**) Design of the abscisic acid (ABA)-inducible molecular interaction recorder. The ABA-responsive interaction domains ABI and PYL1 were fused to the N- and C-terminal fragments of the S7 split monomer, respectively. (**H**) Schematic of ABA-induced interaction between ABI and PYL1, promoting split-monomer reconstitution and subsequent incorporation into the growing protein tape. (**I**) Experimental timeline for ABA-inducible molecular interaction recording in U2OS cells using HaloTag-based timestamps. Cells were treated with 100 μM ABA for either 3- or 6-h terminal treatments or a 12-h intermediate treatment during the 66-h recording period, alongside an untreated control. (**J**) Left: representative confocal images and fluorescence intensity profiles for a tape in the untreated control group. Right: population-level signal monomer intensity change from baseline. *n* = 18 tapes from 18 cells. Throughout this figure: center line, mean; dark shaded boundary, s.e.m.; light shaded boundary, s.d. (**K**) Same as **J**, but for the 3 h terminal treatment group. *n* = 17 tapes from 17 cells. (**L**) Same as **J**, but for the 6 h terminal treatment group. *n* = 19 tapes from 19 cells. (**M**) Same as **J**, but for the 12 h intermediate treatment group. *n* = 19 tapes from 19 cells. Scale bars, 5 μm.

We first generated eight split-monomer designs, named as S1–S8, each from splitting the 1POK(E239Y) monomer at one of its eight flexible loops longer than 6 amino acids (**Figure 2B**). We then tested these designs for recording the rapamycin-induced dimerization of FK506-binding protein (FKBP) and the rapamycin-binding domain of mechanistic target of rapamycin (FRB) (**Figures 2A** and **S1**)^11^. We fused FKBP to the C-terminal of the N-terminal fragment, and FRB to the N-terminal of the C-terminal fragment, of each split-monomer design, with a 24-aa flexible linker at every fusion junction and a unique epitope tag on each of the two fragments for immunofluorescence readout.

Five of the eight designs (S1, S2, S3, S6, and S7) showed clear expression of both split components on CytoTape-vivo tapes^10^ in U2OS cells. All five designs further exhibited increased density towards the tape ends following treatment with 100 nM rapamycin for 2 h, compared to pre-treatment baseline at the tape center (**Figures 2C**, **2D**, **S4A** and **S4B**). We speculated that the sustained signal increase after rapamycin withdrawal may arise from the intracellular retention of rapamycin over days^12^. Importantly, intensity profiles of the N- and C-terminal components were highly correlated across all five candidates, consistent with the split–reconstitution design (**Figure S4C**). By contrast, when either fragment was expressed alone, little or no signal was detected on the tape, indicating that the presence of both split-monomer components is required for tape incorporation (**Figure S3**). Interestingly, the five designs displayed distinct levels of background signal on the tape from spontaneous reconstitution without rapamycin, with S6 having the highest and S7 having the lowest (**Figure S2**). S7 also produced the largest signal-to-background ratio between post- and pre-rapamycin treatment (**Figures 2D** and **S4A**), indicating that S7, referred to as “split-1POK” hereafter, minimizes spontaneous reconstitution while preserving efficient interaction-dependent reconstitution and tape incorporation among the designs (**Figures 2E** and **2F**).

We next asked whether split-monomers could serve as a generalizable recording framework for molecular interactions in live cells beyond that of FKBP, FRB, and rapamycin. To test this, we adapted the abscisic acid (ABA)-dependent protein-protein interaction system^13^, in which the ABA ligand induces the interaction between ABI and PYL1 and can be efficiently washed out from the cell. We fused each of the pair of ABA-responsive interaction domains to the N- and C-terminal fragments of split-1POK, respectively (**Figures 2G**-**2I**). In the absence of ABA, split-1POK signal along the tape remained unchanged (**Figure 2J**). In contrast, ABA treatment for either 3 or 6 h immediately before fixation produced clear increases in the split-1POK signal near the growing tape ends (**Figures 2K** and **2L**). The 6-h treatment produced longer and more intense split-1POK signal increases than those from the 3-h treatment, indicating the split-1POK design can resolve the intensity and duration of signal event across the time axis, a desired capability as a scaffold for future recorder engineering for diverse molecular and cellular events.

We further applied a 12-h ABA treatment and then withdrew it 6 hours before fixation (**Figure 2M**), and observed split-1POK signal first increased then decayed towards the end of the tape, indicating attenuation of split-1POK tape incorporation after ABA withdrawal. Importantly, the split-1POK signal from ABA treatment remained spatially preserved on the tape after ABA withdrawal, implying that the reconstitution of the split-1POK pair was stabilized after tape incorporation. Together, these results establish split-1POK as a versatile interaction-dependent scaffold to sense transient molecular interactions across time in live cells and encode them durably along protein tapes.

### 2.2 Development of TCaMP, a genetically encoded calcium recorder

We next asked whether the split-monomer scaffold could be repurposed as an activity recorder when equipped with protein pairs whose interaction is activity-dependent. We focused on building a recorder for the dynamics of Ca^2+^, a ubiquitous second messenger that governs many intracellular signaling processes (**Figure 3A**)^2^. We selected three pairs of calmodulin (CaM) and its binding peptides (CaMBPs) well-characterized in the literature for Ca^2+^-dependent binding^7,8,14^, and fused CaM or CaMBP to either the N- or C-terminal fragment of split-1POK, yielding six unique pairs of fused fragments as candidate signal monomer designs (**Figure S5A**). Under a 30-min extracellular treatment with 2 μM ionomycin and 5 mM CaCl_2_ to elevate the intracellular Ca^2+^ level (referred to as the “strong stimulation” hereafter) during a 3-day recording period, one design produced a strong increase in signal monomer density, appearing as a sharp, transient response waveform along the tape (**Figures 3B** and **S5B-S5D**). The response showed a median increase of more than 200% relative to the pre-stimulation baseline, with an amplitude approximately 11-fold greater than that observed in the corresponding untreated control group. The magnitude of stimulated signal increase of this design was significantly larger than that of the other five designs, which showed little or no detectable signal increase along the tape under the same condition and whose signal magnitude did not differ significantly from their respective untreated controls (**Figure 3B**). With HaloTag-based timestamping, the onset of the increase in signal monomer density of this design and the onset of dye transition aligned on the tape when Ca^2+^ stimulation and the HaloTag ligand dye switching were initiated simultaneously (**Figures 3E** and **3F**). No corresponding signal increase was observed in unstimulated controls (**Figures S6A and S6B**). Horizontal alignment of individual response waveforms to their respective peak times revealed a median half-rise time (t_1/2_,rise) of 1.84 [1.20, 2.33] h, half-decay time (t_1/2_,decay) of 1.81 [1.00, 3.04] h, and full width at half maximum (FWHM) of 3.70 [2.50, 5.55] h (median [Q1, Q3]) under a strong Ca^2+^ stimulation (**Figure 3L**). We termed this design of “Tape monomer-CalModulin-Peptide” as TCaMP (**Figures 3C** and **3D**).

**Figure 3.**
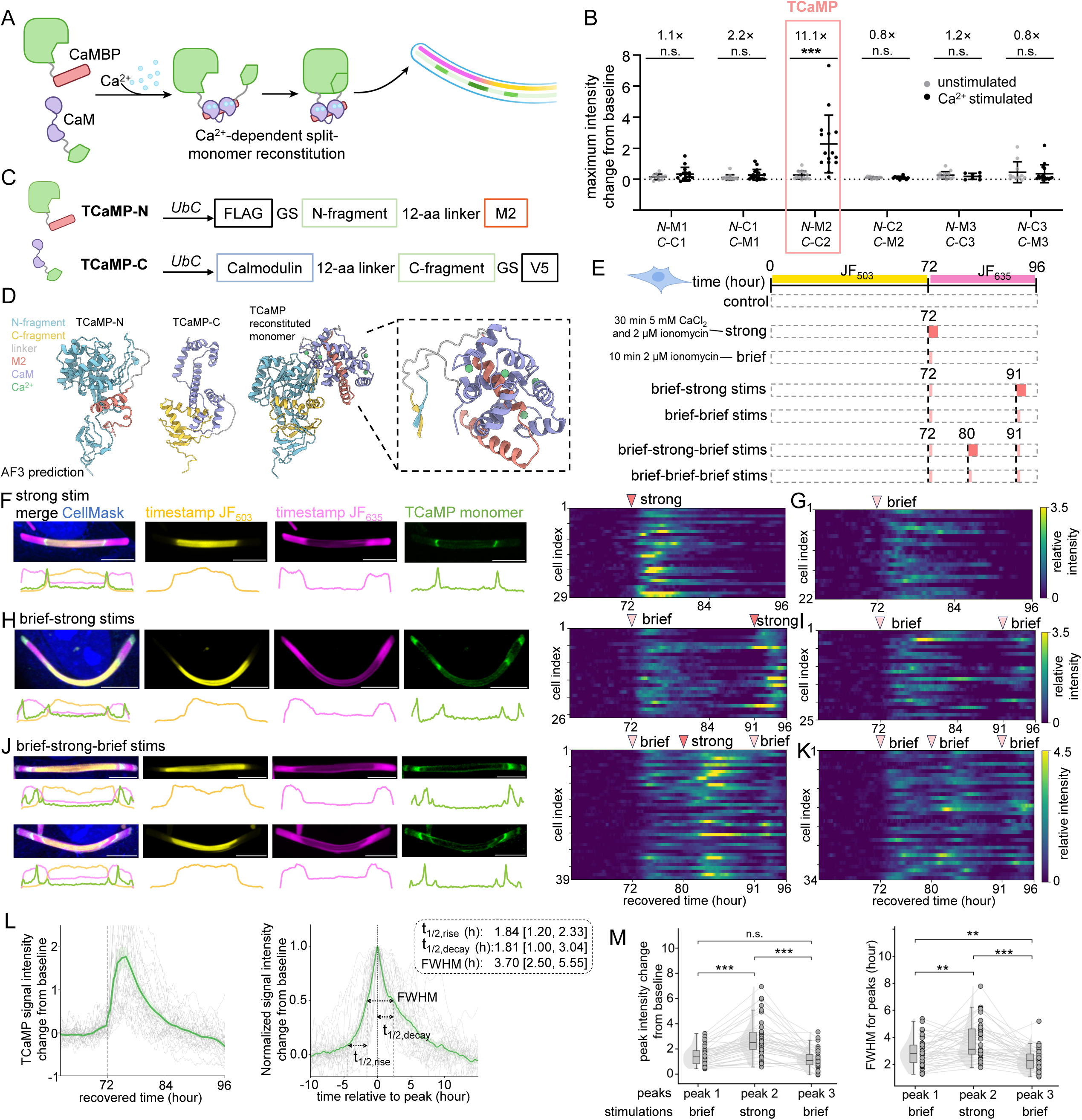
TCaMP, a genetically encoded calcium recorder, records Ca^2+^ dynamics on a single intensity scale across time. (**A**) Schematic of Ca^2+^-dependent split-monomer reconstitution and incorporation into the growing protein tape. Intracellular Ca^2+^ promotes the interaction between calmodulin (CaM) and a CaM-binding peptide (CaMBP), and thus the reconstitution of the split monomer and its subsequent incorporation into the tape. (**B**) Quantification of maximum fluorescence intensity change from baseline in U2OS cells expressing six designs, with (black) or without (gray) Ca^2+^ stimulation (mean ± s.d.). Fold changes of average values between stimulated and unstimulated groups are indicated above each design. The N-M2/C-C2 design, which exhibited the strongest Ca^2+^-dependent response, was selected and named TCaMP (highlighted in red). The numbers of tapes (cells) in the unstimulated/Ca^2+^-stimulated groups were *n* = 11/15, 14/17, 16/15, 12/11, 15/7, and 11/16 for N-M1/C-C1, N-C1/C-M1, N-M2/C-C2, N-C2/C-M2, N-M3/C-C3, and N-C3/C-M3, respectively. Two-sided Mann-Whitney U test, with Holm correction for multiple comparisons. (**C**) Constructs of the TCaMP fragments, TCaMP-N and TCaMP-C. (**D**) AlphaFold 3 (AF3)-predicted structures of TCaMP-N, TCaMP-C, and their reconstituted complex in the presence of four Ca^2+^ ions. The enlarged view (right, dashed box) highlights the Ca^2+^-dependent interaction interface between calmodulin (CaM, purple) and the CaM-binding peptide (M2, red) within the reconstituted complex, including the four Ca^2+^ ions (green). (**E**) Experimental timeline for TCaMP recording of single, double, and triple Ca^2+^ stimulation events within a 24-h window (72–96 h), defined by HaloTag-based timestamps during the overall 96-h experiment. Single strong or brief stimulation was applied at 72 h, double stimulation (brief–strong or brief–brief) at 72 and 91 h, and triple stimulation (brief–strong–brief or brief–brief–brief) at 72, 80, and 91 h. Brief stimulation consisted of 10 min treatment with 2 μM ionomycin in cell culture medium; strong stimulation consisted of 30 min treatment with 2 μM ionomycin supplemented with 5 mM CaCl_2_ in cell culture medium. (**F**) Representative confocal images and corresponding fluorescence intensity profiles of TCaMP tapes (left), and a heatmap of individual TCaMP fluorescence intensity profiles plotted over recovered time from the single strong Ca^2+^ stimulation group (right). Each row represents an individual tape, *n* = 29 tapes from 29 cells. Throughout the figure, arrowheads indicate the timing of Ca^2+^ stimulation, with dark red and light red indicating strong and brief stimulations, respectively. (**G**) Heatmap of individual TCaMP fluorescence intensity profiles plotted over recovered time from the single brief Ca^2+^ stimulation group, *n* = 22 tapes from 22 cells. (**H**) Same as **F**, but for the brief-strong double Ca^2+^ stimulations group. *n* = 26 tapes from 26 cells. (**I**) Heatmap of individual TCaMP fluorescence intensity profiles plotted over recovered time from the brief-brief double Ca^2+^ stimulations group, *n* = 25 tapes from 25 cells. (**J**) Same as **F**, but for the brief-strong-brief triple Ca^2+^ stimulations group. *n* = 39 tapes from 39 cells. (**K**) Heatmap of individual TCaMP fluorescence intensity profiles plotted over recovered time from the brief-brief-brief triple Ca^2+^ stimulations group, *n* = 34 tapes from 34 cells. (**L**) Characterization of TCaMP fluorescence response kinetics using the same single strong Ca^2+^ stimulation dataset as in **F**. Left: TCaMP fluorescence intensity changes from baseline plotted over recovered time, with individual traces aligned to the stimulation onset at 72 h. Right: the same traces aligned to their individual peak times (time = 0 h) and normalized to their respective baseline and peak intensities. The half-rise time (t_1/2,rise_) was defined as the time interval from baseline to half-maximum intensity on the rising phase; the half-decay time (t_1/2,decay_) was defined as the time interval from peak intensity to half-maximum intensity on the decay phase; full width at half maximum (FWHM) was defined as the time interval between the half-maximum intensity crossings on the rising and decay phases. Median values and interquartile ranges (IQRs) for each parameter are indicated in the corresponding plots. *n* = 29 tapes from 29 cells; center line, mean; dark shaded boundary, s.e.m.; light shaded boundary, s.d.; gray thin lines, data from individual tapes. (**M**) Quantification of peak fluorescence intensity change from baseline (left) and full width at half maximum (FWHM; right) for the three sequential TCaMP responses recorded in **J**. Raincloud plots show data distributions, with half-violins representing density estimates, boxes indicating medians and IQRs, and whiskers extending to 1.5 × IQR. Gray dots represent individual tapes, with lines connecting measurements from the same tape across the three peaks. *n* = 39 tapes from 39 cells; paired Wilcoxon signed-rank tests, with Holm correction for multiple comparisons. Throughout this figure: n.s., not significant; **, P < 0.01; ***, P < 0.001. Scale bars, 4 μm.

To further characterize TCaMP for temporal recording of intracellular calcium dynamics, we performed a series of Ca^2+^ stimulation experiments varying both stimulus strength and timing. TCaMP responded robustly to a 10-min 2 μM ionomycin-only treatment, hereafter referred to as the “brief stimulation”, with a 163% median increase from pre-stimulation baseline (**Figures 3E**, **3G** and **S6C**). We further quantified the TCaMP response amplitude under brief and strong stimulations (**Figure S6F**). We next examined whether TCaMP could resolve multiple transient Ca^2+^ events within the same recording period (**Figure 3E**). When cells were stimulated twice with a 19-h interval, TCaMP reproducibly generated two distinct signal responses along the tape (**Figures 3H**, **3I** and **S6D**). Moreover, when the second stimulation differed in strength, the strong-stimulation group exhibited a significantly greater second-response amplitude than the brief-stimulation group (**Figure S6G**). We further tested whether TCaMP could resolve three sequential Ca^2+^ events spaced within a 24-h period (**Figure 3E**). In a paradigm consisting of a brief–strong–brief stimulation sequence, three distinct responses were reproducibly detected along the tape (**Figure 3J**). The second response, corresponding to the strong stimulation, exhibited significantly larger peak amplitude and full width at half maximum (FWHM) than those of the first and third responses in paired comparisons (**Figure 3M**). By contrast, when all three stimulations were brief, the three recorded responses showed more homogeneous amplitudes and widths, with the final response showing a modest reduction in amplitude relative to the second response (**Figures 3K, S6E, S6H** and **S6I**).

### 2.3 TCaMP records endogenous calcium dynamics from receptor activation and intracellular store release

In addition to its roles in intracellular processes, cytosolic Ca^2+^ is often used to report events beyond the cytosol, such as the activity of upstream receptors at the plasma membrane and the handling of Ca^2+^ in organelles^15,16^. For example, Ca^2+^ mobilization is a common functional readout of GPCR activation^17^, widely used in receptor characterization and ligand screening in drug discovery^16^. Cytosolic Ca^2+^ is also commonly used as a readout to study Ca^2+^ release from endoplasmic reticulum (ER) stores and its regulation^18^. To demonstrate TCaMP recording of Ca^2+^ dynamics in these contexts, we applied two widely used pharmacological agents, histamine and thapsigargin (**Figures 4A** and **4B**). Specifically, histamine activates the Gq-coupled H1 receptor to release Ca^2+^ from ER stores, producing transient Ca^2+^ elevations^19^.

**Figure 4.**
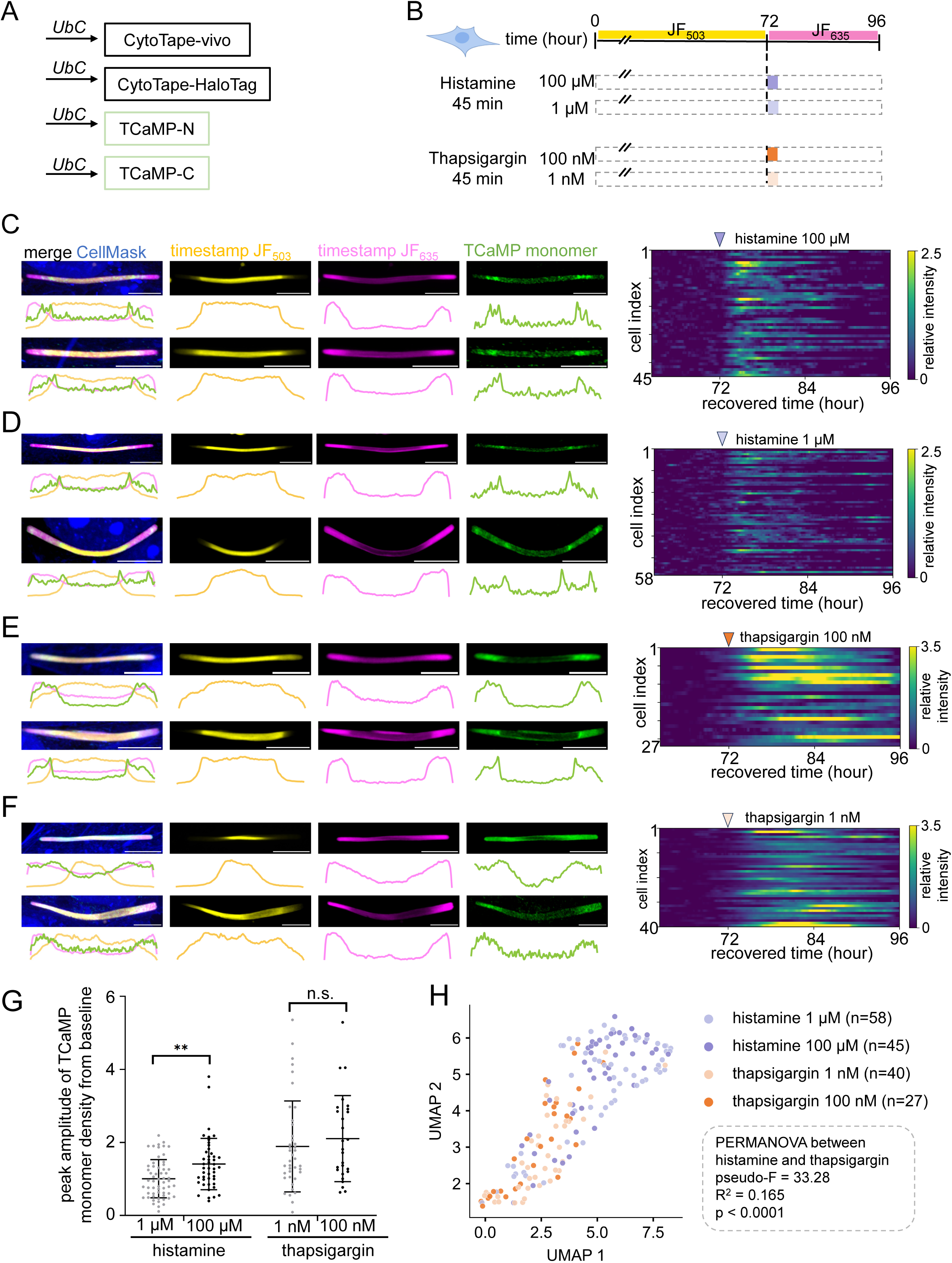
TCaMP records Ca^2+^ dynamics from GPCR activation and intracellular store release. **(A)** TCaMP constructs were co-expressed with CytoTape-vivo and a timestamp monomer under the *UbC* promoter in U2OS cells. (**B**) Experimental timeline for TCaMP recording of drug-induced endogenous Ca^2+^ signaling dynamics within a 24-h window (72–96 h), defined by HaloTag-based timestamps during the overall 96-h experiment. Stimulations with different concentrations of histamine or thapsigargin were applied at 72 h for 45 min. (**C**) Representative confocal images and corresponding fluorescence intensity profiles of TCaMP tapes (left), and a heatmap (right) of individual TCaMP fluorescence intensity profiles plotted over recovered time from the 100 μM histamine stimulation group. Each row represents an individual tape, *n* = 45 tapes from 45 cells. Arrowheads mark stimulation onset, with darker and lighter shades indicating higher and lower drug concentrations, respectively; violet denotes histamine, and orange denotes thapsigargin. Throughout this figure, scale bars, 5 μm. (**D**) Same as **C**, but for the 1 μM histamine stimulation group. *n* = 58 tapes from 58 cells. (**E**) Same as **C**, but for the 100 nM thapsigargin stimulation group. *n* = 27 tapes from 27 cells. (**F**) Same as **C**, but for the 1 nM thapsigargin stimulation group. *n* = 40 tapes from 40 cells. (**G**) Quantification of peak TCaMP monomer density changes from baseline in response to different concentrations of histamine or thapsigargin (mean ± s.d.). *n* = 58, 45, 40, and 27 tapes from 58, 45, 40, and 27 cells for 1 μM histamine, 100 μM histamine, 1 nM thapsigargin, and 100 nM thapsigargin, respectively. n.s., not significant; **P < 0.01; ***P < 0.001; two-sided Mann–Whitney U tests with Holm correction for multiple comparisons. (**H**) UMAP visualization of single-cell TCaMP fluorescence trajectory shapes over the 24-h recording window following stimulation with histamine (1 μM, light violet, *n* = 58 cells; 100 μM, dark violet, *n* = 45 cells) or thapsigargin (1 nM, light orange, *n* = 40 cells; 100 nM, dark orange, *n* = 27 cells). Each signal trajectory was z-score normalized before PCA–UMAP analysis. Drug groups were compared with concentrations pooled, using PERMANOVA on Euclidean distances between the normalized trajectories (pseudo-*F* = 33.28, *R*^2^ = 0.165, *p* < 0.0001; cell-label permutations). Same dataset as **C**–**F**.

Thapsigargin irreversibly inhibits the sarco/endoplasmic reticulum Ca^2+^-ATPase (SERCA), depleting ER stores and preventing their refilling, resulting in a slower rise in cytosolic Ca^2+^ that persists well beyond the treatment period^20^. Following a 45-min stimulation with either 1 μM or 100 μM histamine, many TCaMP-expressing cells exhibited clear, transient responses in signal monomer density along the tape (**Figures 4C** and **4D**). In addition, some cells displayed multi-peak or oscillatory waveforms in the following recording window after the initial Ca^2+^ response. These hour-scale waveform features may reflect prolonged or recurrent Ca^2+^ activity following histamine stimulation rather than fast Ca^2+^ oscillations at minute-scale frequencies. These observations highlight TCaMP’s unique capability to scalably record hour-scale temporal features across days, at the cost of slower kinetics compared to real-time live-cell imaging of fluorescent calcium indicators. Across the population, the overall Ca^2+^ signal amplitude was greater following 100 μM histamine than that following 1 μM histamine, consistent with previous reports showing that stronger histamine stimulation recruits a larger fraction of responding cells and evokes larger Ca^2+^ responses (**Figure 4G**)^21^. After 1 nM or 100 nM thapsigargin treatment, in contrast, most cells under both conditions exhibited a slow rise and prolonged elevation of the recorded TCaMP signal, with peak amplitudes showing no detectable difference between the two concentrations (**Figures 4E**-**4G**). These results are in alignment with the pharmacological feature of thapsigargin exhibiting sub-nanomolar affinity and prolonged effect in blocking SERCA and increasing cytosolic Ca^2+^ in the 24 hours following treatment^22^. Principal component analysis followed by a UMAP visualization (**Figure 4H**) revealed distinct temporal features of the response waveforms between histamine and thapsigargin treatment groups (**Figures 4C**-**4F**). Together, these results indicate that TCaMP records calcium dynamics not only from ionophore-mediated transmembrane entry but also from GPCR activation and from intracellular store release, demonstrating a genetically encoded calcium recorder that resolves signal amplitude in analog and temporal signature continuously across time.

## 3 Discussion

In this work, we established activity-dependent split-monomer reconstitution as a programmable strategy for molecular recording of cellular activities and molecular dynamics. We identified split-1POK pairs that efficiently reconstitute into complete monomers upon molecular interaction of their fused domains. By coupling split-monomer reconstitution to intracellular calcium and molecular interactions, we developed TCaMP, a genetically encoded calcium recorder, as well as ligand-dependent molecular interaction recorders. In each case, sensing and writing are carried out by the same tape monomers already present in the cytosol, decoupling recording from transcription- or translation-associated cascades.

The recording kinetics of these systems are governed by several interconnected processes. The ON kinetics primarily involve two sequential steps: the activity of interest induces the interaction between domains of interest and subsequent reconstitution of split monomers, followed by incorporation of the reconstituted monomers into the elongating protein tape. In contrast, the OFF kinetics reflect two distinct, possible outcomes of the reconstituted monomers: incorporation into the protein tape from the cytosolic monomer pool, or dissociation back into individual split monomers before tape incorporation. Before reaching these outcomes, reconstituted monomers may diffuse in the cytosolic pool and continue to be incorporated into the tape even after the initiating stimulus has subsided. In U2OS cells, TCaMP exhibited a median half-rise time of 1.84 h and a median half-decay time of 1.81 h under a 30 min stimulation with 2 μM ionomycin and 5 mM CaCl_2_. We envision that the temporal resolution of TCaMP is fundamentally dependent on the physical processes underlying protein-tape recording, including the intrinsic tape growth rate, the kinetics of monomer incorporation from the cytosolic pool, and the stimulus-dependent association and dissociation of split monomers. Improving these kinetics therefore represents an important direction for future development, requiring systematic characterization and engineering of three key components: the split-monomer pairs, the protein-tape scaffold, and the calcium-sensitive interaction domains. Advances in AI-guided protein engineering and binder design may provide promising strategies for optimizing these components^23^. In parallel, improving the temporal accuracy of retrospective signal reconstruction will require faster and potentially multi-round molecular timestamping methods. The split-tape monomer strategy itself may provide a versatile foundation for developing such methods through chemically or optically inducible dimerization^24,25^.

TCaMP resolves intracellular calcium dynamics with hour-scale kinetics over days, complementing fluorescent calcium indicators that offer sub-second kinetics under real-time live-cell imaging, by trading temporal resolution for scalability in applications that favor long-term, large-scale calcium recording. TCaMP also provides an approach distinct from calcium integrators and calcium-dependent cell labeling methods by recording calcium signal intensities on the same intensity scale across a continuous time axis within individual cells, providing quantitative comparison of calcium levels over time within the same cell. Besides tracking cytosolic calcium dynamics, TCaMP can also be applied to study processes beyond the cytosol, including transmembrane calcium entry, receptor activity, and intracellular store release, by recording their calcium signatures across time. Together, TCaMP and other split-monomer-based recorders offer immediate utility for studying large-scale, long-term cellular processes, even as the framework itself retains significant potential for future growth.

## 4 Methods

### Mammalian cell line

Human bone osteosarcoma epithelial cells (referred to as “U2OS” throughout this paper; ATCC HTB-96) were grown in Dulbecco’s Modified Eagle’s Medium (DMEM; Gibco; containing high glucose, GlutaMAX supplement, and pyruvate) supplemented with 10% HI FBS (Corning) and penicillin/streptomycin (Gibco), referred to as the “U2OS culture medium” throughout this paper. Cells were grown in a cell culture incubator at 37 °C with 5% CO_2_.

### Molecular cloning

Genes were synthesized by Twist Bioscience, and then inserted into mammalian expression vectors, pAAV-*UbC*, from our previous work^10^, via Gibson assembly. See **Table S1** for the sequences of genes and motifs in this study.

### Plasmid transfection in U2OS cells

U2OS cells at a low passage number (<10 passages) were plated at 40% confluence onto a 100-mm tissue culture dish. Cells were then grown in the U2OS culture medium in a cell culture incubator at 37 ^◦^C with 5% CO_2_. After reaching 50-70% confluence, cells were transferred to 24-well glass-bottom plates via trypsin treatment. The 24-well glass-bottom plates were pre-treated with 70 µL Matrigel in the cell culture incubator for 40 min before cell plating. 24 h after cell plating, genes were delivered using the *Trans*IT-X2 Dynamic Delivery System kit (Mirus Bio). For split-1POK engineering experiments (**Figures 2A**-**2F**), each well of a 24-well glass-bottom plate received 100 ng structural monomer plasmid (*UbC*-CytoTape-vivo-HA), 20 ng timestamp monomer plasmid (*UbC*-CytoTape-HaloTag), 10 ng of N-fragment plasmid from each design, 10 ng of C-fragment plasmid from each design, and 160 ng pUC19 plasmid. For single-fragment control experiments in split-1POK engineering (**Figure S3**), each well of a 24-well glass-bottom plate received 100 ng structural monomer plasmid (*UbC*-CytoTape-vivo-HA), 20 ng timestamp monomer plasmid (*UbC*-CytoTape-HaloTag), 10 ng of either N- or C-fragment plasmid from each design, and 170 ng pUC19 plasmid. For experiments related to ABA-inducible molecular interaction recording (**Figures 2G**-**2M**), each well of a 24-well glass-bottom plate received 100 ng structural monomer plasmid (*UbC*-CytoTape-vivo-HA), 20 ng timestamp monomer plasmid (*UbC*-CytoTape-HaloTag), 10 ng of N-fragment design (*UbC*-FLAG-N_fragment-ABI), 10 ng of C-fragment design (*UbC*-PYL1-C_fragment-V5), and 160 ng pUC19 plasmid. For experiments related to screening of Ca^2+^-responsive split-monomer designs (**Figure 3B**), each well of a 24-well glass-bottom plate received 100 ng structural monomer plasmid (*UbC*-CytoTape-vivo-HA), 20 ng timestamp monomer plasmid (*UbC*-CytoTape-HaloTag), 10 ng of N-fragment plasmid from each design, 10 ng of C-fragment plasmid from each design, and 160 ng pUC19 plasmid. For experiments related to TCaMP recording (**Figures 3F**-**3K, 4C**-**4F**), each well of a 24-well glass-bottom plate received 100 ng structural monomer plasmid (*UbC*-CytoTape-vivo-HA), 20 ng timestamp monomer plasmid (*UbC*-CytoTape-HaloTag), 10 ng of TCaMP-N (*UbC*-FLAG-N_fragment-M2), 10 ng of TCaMP-C (*UbC*-Calmodulin-C_fragment-V5), and 160 ng pUC19 plasmid. Plasmids were first diluted in 50 µL Opti-MEM medium, followed by the addition of 0.9 µL *Trans*IT-X2 reagent. The mixture was incubated at room temperature for 25 min, and then added to every single well.

### Timestamping via HaloTag ligand dyes

HaloTag-based timestamp was implemented with cell-permeant HaloTag ligand Janelia Fluor (JF) dyes (Promega) (**Figure 3** and related figures; **Figure 4** and related figures). JF_503_ and JF_635_ were used in this study for timestamps. The JF dyes were in lyophilized powder form and stored at -20 °C before use. The JF dye powder was first dissolved in 50 µL dimethyl sulfoxide (DMSO) and then diluted into 10 mL of U2OS cell culture medium at a final concentration of 0.1 µM. The dyed medium was filtered using a sterile 0.22 µm syringe filter before being added to the culture. It was then used to replace the original medium in cell cultures at designated time points. During the dye-switching process, the medium containing the original dye was fully removed, followed by thorough washes of the cell cultures three times with 37 °C DPBS. Finally, the cells in each well of the 24-well plate were cultured in 500 µL of fresh U2OS cell culture medium supplemented with a new JF dye at 0.1 µM final concentration in the cell culture incubator at 37 °C with 5% CO_2_.

### Rapamycin treatment in U2OS

Rapamycin (MedChemExpress HY-10219) was stored as a 10 mM stock solution dissolved in DMSO at –80 °C before use. Before treatment, the 10 mM rapamycin stock was diluted into U2OS culture medium to a final concentration of 100 nM. The original culture medium was removed, and 500 μL of rapamycin-containing medium was added to each well. Cells were treated for durations as indicated in the Results section. After treatment, the drug-containing medium was removed, and cells were washed three times with warmed DPBS before fresh U2OS culture medium was added and cells were returned to the incubator.

### Abscisic acid treatment in U2OS

Abscisic acid (ABA, MedChemExpress HY-100560) was stored as a 10 mM stock solution dissolved in DMSO at –20 °C before use. Before treatment, the 10 mM ABA stock was diluted into U2OS culture medium to a final concentration of 100 μM. The original culture medium was removed, and 500 μL of ABA-containing medium was added to each well. Cells were treated for durations as indicated in the Results section. After treatment, the drug-containing medium was removed, and cells were washed three times with warmed DPBS before fresh U2OS culture medium was added and cells were returned to the incubator.

### Ionomycin, CaCl_2_, histamine, and thapsigargin treatments in U2OS

Ionomycin (Invitrogen, I24222) was prepared as a 1 mM stock solution in DMSO and stored at −20 °C until use. Calcium chloride dihydrate (CaCl_2_·2H_2_O) was dissolved in water to prepare a 1 M CaCl_2_ stock solution. Before treatment, the stock solutions were diluted in the U2OS culture medium to final concentrations of 2 μM ionomycin and 5 mM CaCl_2_ for strong Ca^2+^ stimulation, or 2 μM ionomycin alone for brief Ca^2+^ stimulation. For histamine stimulation of TCaMP, histamine (Millipore Sigma, H7125) was prepared as a 500 mM stock solution in water and stored at −20 °C until use. Before treatment, the 500 mM histamine stock was diluted into U2OS culture medium to a final concentration of 100 μM or 1 μM. For thapsigargin treatment, thapsigargin (MedChemExpress, HY-13433) was prepared as a 1 mM stock solution in DMSO and stored at −80 °C until use. Before treatment, the 1 mM thapsigargin stock was diluted into U2OS culture medium to a final concentration of 100 nM or 1 nM. The original culture medium was removed, and 500 μL of stimulation medium was added to each well. Cells were treated for durations as indicated in the Results section. After treatment, the drug-containing medium was removed, and cells were washed three times with warmed DPBS before fresh U2OS culture medium was added and cells were returned to the incubator.

### Antibodies and cell morphology stains

Primary antibodies (1:1000 dilution): anti-V5 (Invitrogen R960-25), anti-FLAG (Invitrogen 740001), anti-HA (Cell Signaling Technology 3724; Invitrogen 26183). Fluorescent secondary antibodies (1:1000 dilution): Goat anti-Mouse IgG1 Alexa Fluor 488 (Invitrogen A-21127), Goat anti-Mouse IgG2a Alexa Fluor 488 (Invitrogen A-21131), Goat anti-Mouse IgG2a Alexa Fluor 546 (Invitrogen A-21133), Goat anti-Rabbit IgG (H+L) Alexa Fluor Plus 488 (Invitrogen A-32731), Goat anti-Rabbit IgG (H+L) Alexa Fluor 647 (Invitrogen A-21245), and Goat anti-Mouse IgG1 Alexa Fluor 647 (Invitrogen A-21240). Cell morphology stains (1:1000 dilution): HCS CellMask™ Stains Blue (Invitrogen, H32720), NeuroTrace 435/455 Blue Fluorescent Nissl Stain (Invitrogen N21479).

### Immunostaining of epitope tags and cell morphology staining

U2OS cells were fixed in 10% buffered formalin (Fisher Scientific) at room temperature (RT) for 10 min, followed by three 5-minute washes in 1×PBS at RT. Blocking was performed in MAXblock blocking medium (Active Motif) supplemented with 0.1% Triton X-100 and 100 mM glycine for 20 min at RT, followed by three additional 5-min washes in MAXwash washing medium (Active Motif) at RT. Cells were then incubated with primary antibodies diluted in MAXbind staining medium (Active Motif) for 1 h at RT. Afterward, cells underwent three 5-min washes in MAXwash washing medium at RT. Secondary antibodies were applied in MAXbind staining medium and incubated for 1 h at RT. Cells were then washed three times with MAXwash washing medium for 5 min each at RT. Finally, cells were incubated with HCS CellMask™ Blue Stain or NeuroTrace 435/455 Blue Fluorescent Nissl Stain (Invitrogen) for 30 min and stored in 1×PBS at 4 °C until imaging.

### Fluorescence microscopy of immunostained cells

Fluorescence microscopy was performed using a spinning disk confocal microscope (Yokogawa CSU-W1 Confocal Scanner Unit on a Nikon Eclipse Ti2 inverted microscope) equipped with a 10× objective, a 40× objective, and a 100× objective, a Hamamatsu ORCA-Fusion BT sCMOS camera controlled by the NIS-Elements AR software, and laser/filter sets for 405 nm, 488 nm, 561 nm, and 640 nm optical channels. Multi-channel volumetric imaging was conducted at 0.4 µm per Z-step for each field of view under the 40× objective. Imaging parameters remained consistent across all samples within each experimental set.

### Software for image analysis

Image analysis was performed in Python and ImageJ (National Institutes of Health).

### Readout of intensity profile along the tape

Fluorescence intensity profiles along the tapes were extracted from images as previously described^9^.

### Time recovery from timestamps

Two HaloTag ligand dyes were used per experiment in this study to recover time axes in U2OS cells (**Figures 3F-3K** and related figures; **Figures 4B-4F**). The methods for identifying the onset of JF dye switching along the tape (“timestamp”) and recovering the time axis from timestamps along the tape were performed as previously described^10^.

### Protein structure prediction and visualization

To visualize the N- and C-terminal fragments of split-1POK (**Figures 2F** and **S1**), the corresponding split sites were mapped onto the experimentally determined X-ray crystal structure of 1POK (PDB ID: 1POK^26^). For regions unresolved in the crystal structure (**Figure 2F**), structural predictions were generated using AlphaFold 3 (AF3) through the AlphaFold Server^27^. The structures of TCaMP-N and TCaMP-C, as well as their reconstituted complex in the presence of Ca^2+^ (**Figure 3D**), were also predicted using AF3 through the AlphaFold Server. All protein structures were visualized using UCSF ChimeraX^28^.

### Statistical analysis

Statistical analyses were performed using GraphPad Prism and Python. All statistical tests were two-sided where applicable, except for the upper-tail permutation test in **Figures 4H and S4B**.

## Acknowledgments

We thank Jiahui Ding, Ilgang Hong, Yongjie Hou, Bianca Wang, Wenjing Wang, Nan Xiang, and Kairui Yang for discussions and technical assistance. Y.Y. acknowledges the Rackham Graduate Student Research Grant. C.L. is supported by the National Institutes of Health (NIH) Director’s New Innovator Award (DP2MH140133), the NIH/NINDS (R01NS144823), the Glenn Foundation for Medical Research and American Federation for Aging Research Grant Award for Junior Faculty, and the Klingenstein Fellowship Award in Neuroscience.

## Author contributions

Y.Y. and C.L. conceived the concept of the molecular strategies and developed the overall research plan. Y.Y. designed the constructs, performed experiments, and analyzed the data. Y.Y. and C.L. interpreted the data and wrote the manuscript. C.L. supervised the project.

## Data availability

Sequences of genes and motifs in this study are available in **Table S1**. Plasmids will be available at Addgene upon completion of the deposition process.

---

**Figure S1.**
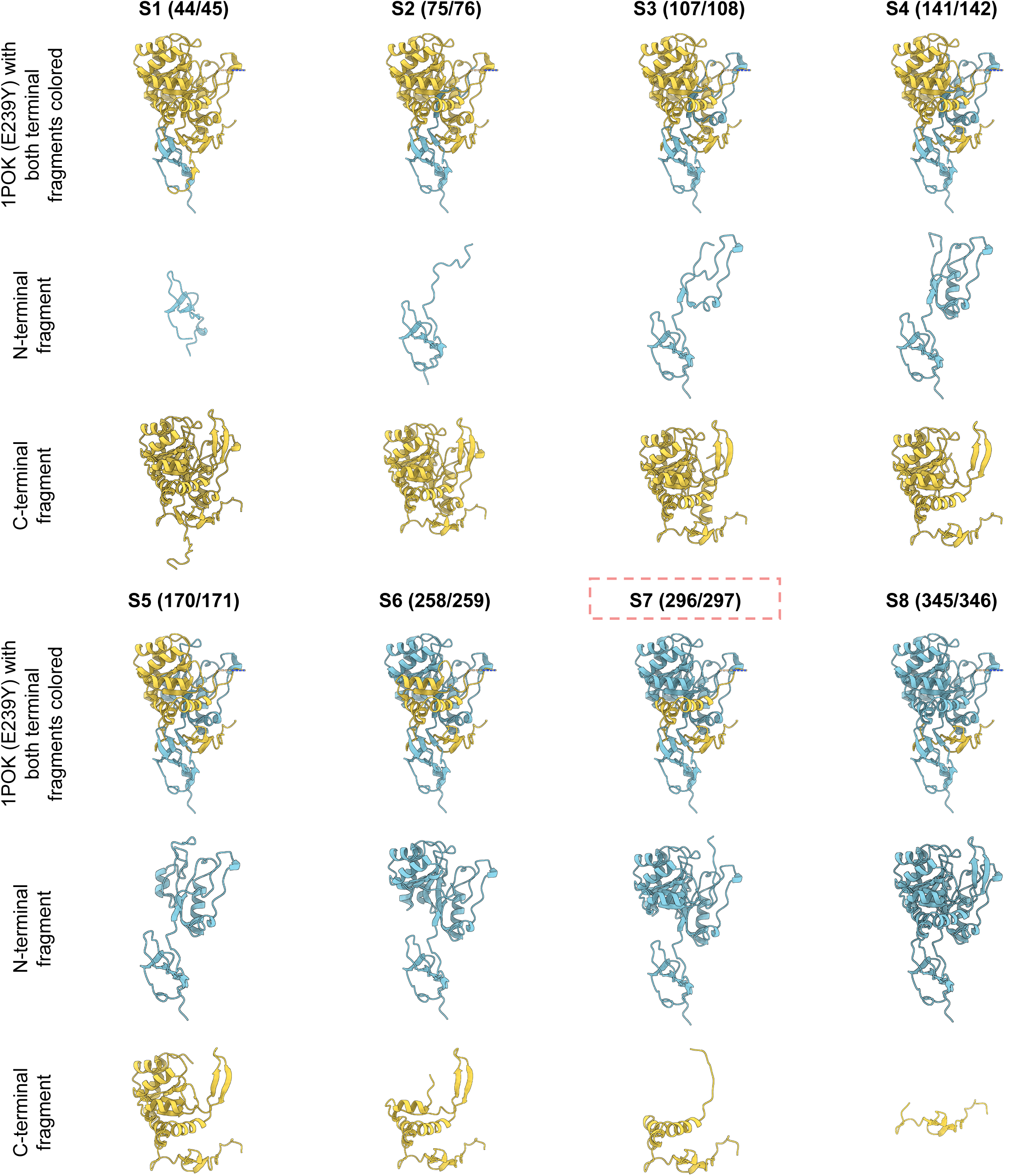
Structural representations of eight split-1POK designs. Structural representations of the eight engineered split sites (S1–S8) in 1POK(E239Y). For each split site, the top panel shows the crystal structure of PDB: 1POK, with regions corresponding to the N- and C-terminal fragments colored in blue and yellow, respectively. The middle and lower panels show the corresponding N- and C-terminal fragments of 1POK(E239Y), respectively, with their structures predicted using AlphaFold 3. Amino acid positions defining each split site are indicated above the structures. The design with split site S7 (296/297) is selected as the final split-1POK design and is highlighted by a red dashed rectangle.

**Figure S2.**
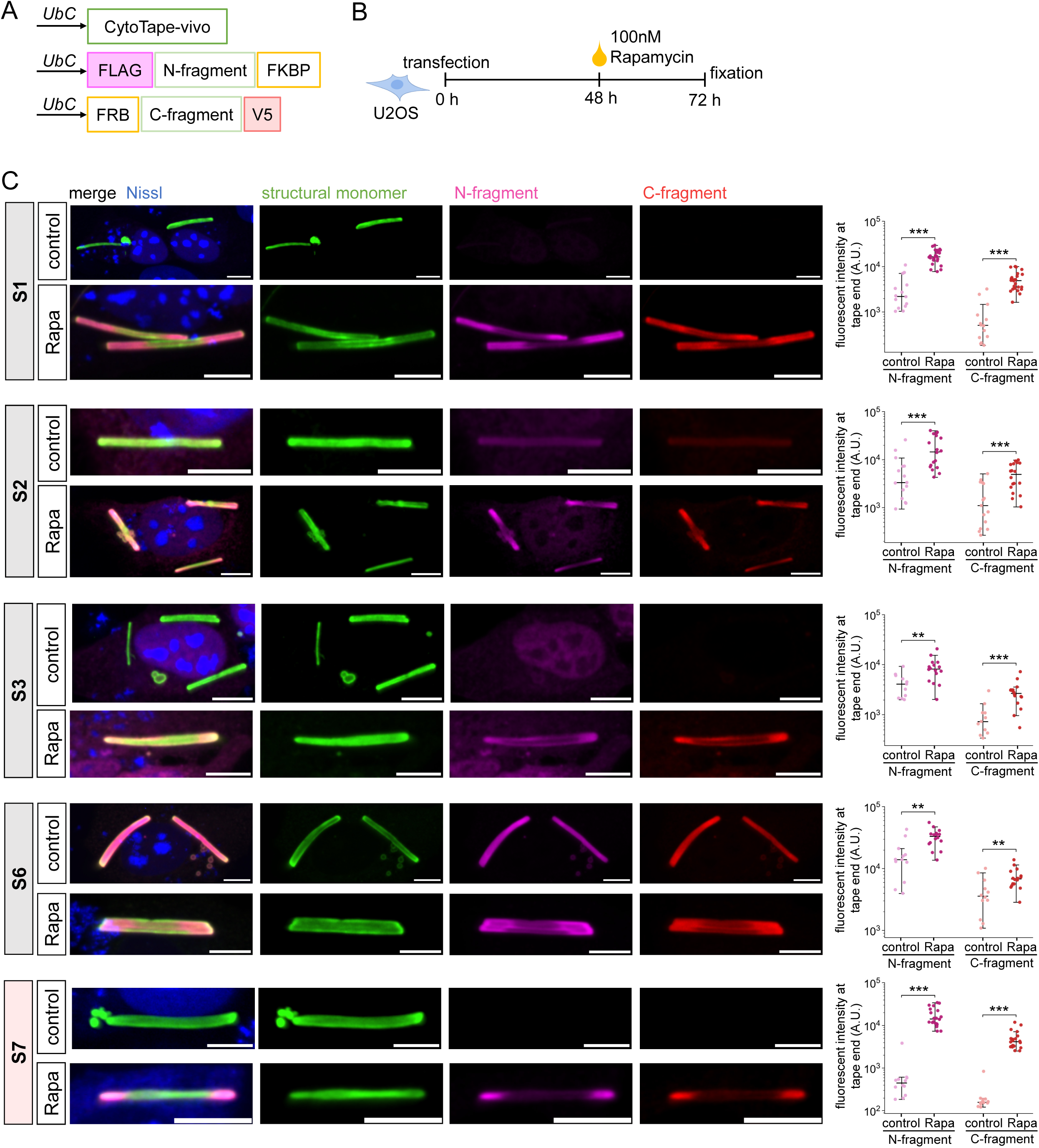
Rapamycin-induced reconstitution of split-1POK designs in U2OS cells. (**A**) Design of the rapamycin-inducible split-monomer recorder. The rapamycin-responsive interaction domains FKBP and FRB were fused to the N- and C-terminal fragments of the split monomer, respectively, with FLAG and V5 epitope tags for fluorescence detection. Both constructs were co-expressed with CytoTape-vivo under the *UbC* promoter. (**B**) Experimental timeline for rapamycin-inducible split-monomer recording in U2OS cells. Cells were transfected at 0 h, treated with 100 nM rapamycin (Rapa) at 48 h, and fixed at 72 h. Same condition as in **Figures 2C**-**2E**. (**C**) Representative confocal images of CytoTape-vivo tapes co-transfected with five split-monomer designs (S1, S2, S3, S6, and S7), with or without Rapa treatment. Corresponding scatter plots (right) quantify raw fluorescence intensity at tape ends for the N- and C-terminal fragments in control (light magenta/red) and Rapa-treated (dark magenta/red) groups. Dots represent individual tapes, horizontal bars indicate medians, and whiskers extend to the most extreme values within 1.5 × IQR. The selected split site, S7, is highlighted in red. **P < 0.01; ***P < 0.001; two-sided Mann–Whitney U tests. Scale bars, 10 μm.

**Figure S3.**
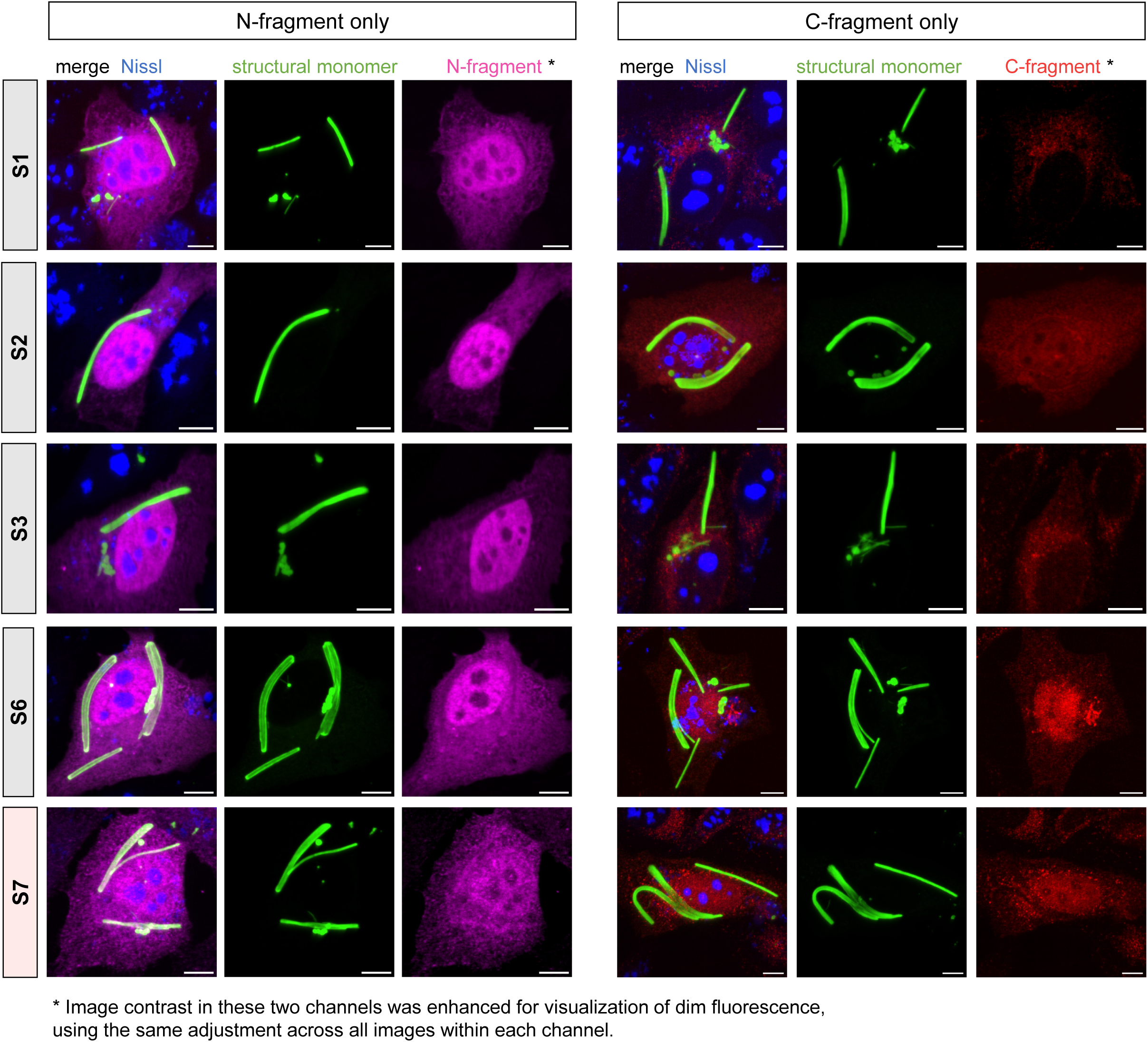
A single N- or C-fragment from split-1POK designs alone exhibits little or no detectable incorporation into protein tapes. Representative confocal images of U2OS cells expressing the five split-monomer designs (S1, S2, S3, S6, and S7) fused to FKBP and FRB as in **Figure S2**, but with only one of the paired N- or C-terminal fragment constructs co-transfected. Images show merged channels with Nissl staining (blue), structural monomer (green), and either the N-terminal fragment (magenta, left) or C-terminal fragment (red, right). The selected split site, S7, is highlighted in light red. Contrast settings were kept consistent within each fragment channel (N or C) and adjusted to visualize weak background fluorescence. Scale bars, 10 μm.

**Figure S4.**
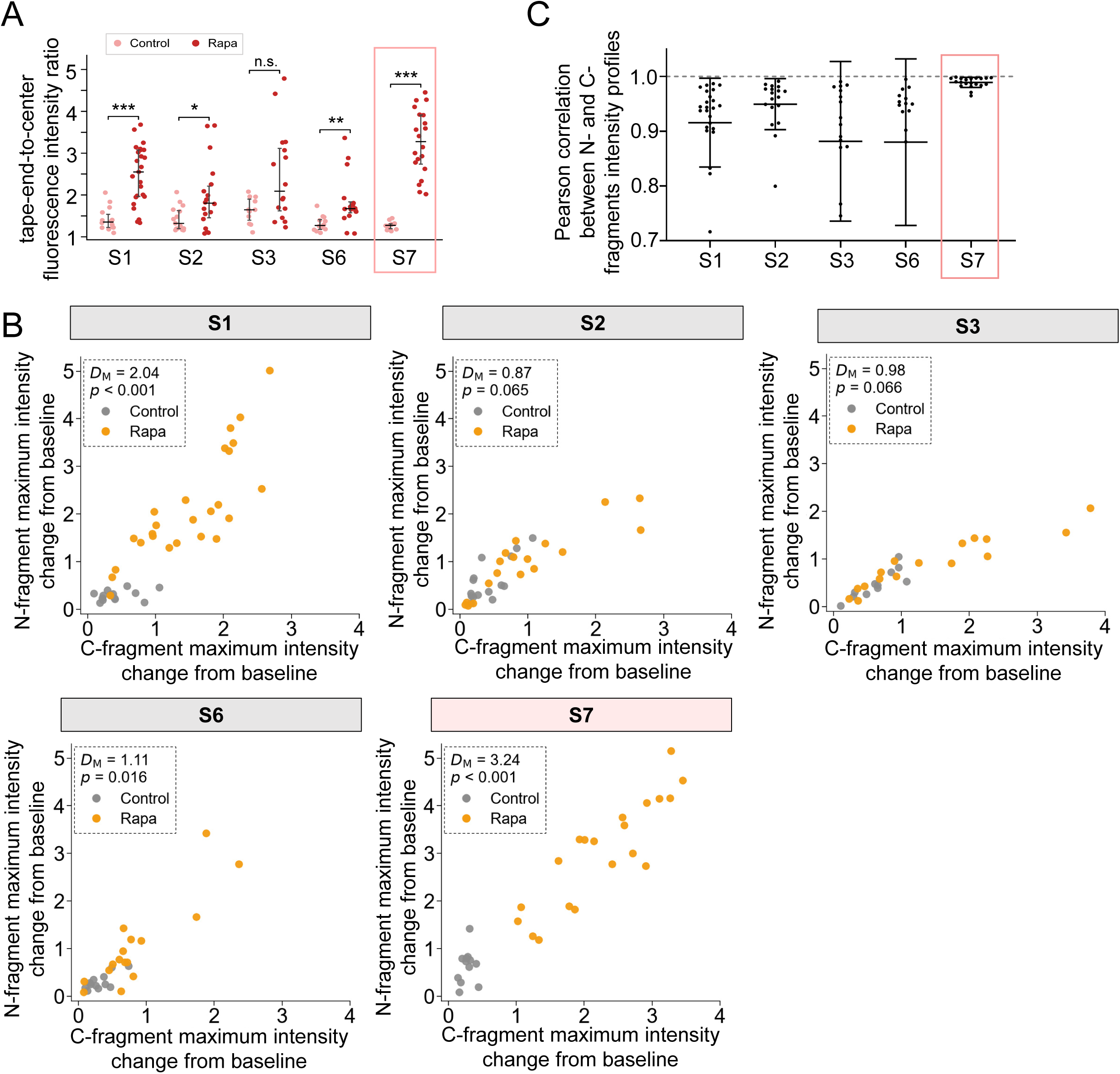
Additional characterizations of split-1POK designs fused to FKBP and FRB under rapamycin-induced reconstitution. (**A**) Quantification of tape end-to-center fluorescence intensity ratio for the five split-monomer designs fused to FKBP and FRB as in **Figure S2**, with (darker red) or without (lighter red) rapamycin (Rapa) treatment (median with IQR), measured using the epitope tag fused to the C-terminal fragment. The tape/cell numbers in the control/Rapa groups are *n* = 14/25, 15/18, 11/16, 12/16, 12/20 for S1, S2, S3, S6, S7 respectively. n.s., not significant; *, P < 0.05; **, P < 0.01; ***, P < 0.001; two-sided Mann-Whitney U test, with Holm correction for multiple comparisons. (**B**) Scatter plots of the maximum fluorescence intensity change from baseline for the N- and C-terminal fragments for control (gray) and Rapa (orange) groups in five split-monomer designs exhibiting detectable expression. Mahalanobis distance (D_M_) between the control and Rapa groups was calculated using their mean vectors and pooled covariance matrix. Statistical significance was assessed using a permutation test with 10,000 random shuffles of group labels, with *p* values calculated as the proportion of permuted distances greater than or equal to the observed distance. The tapes (cells) number in the control/Rapa groups are *n* = 14/25, 15/18, 11/16, 12/16, 12/20 for S1, S2, S3, S6, S7 respectively. (**C**) Quantification of the Pearson correlation between the N- and C-fragments intensity profiles for five split-monomer designs exhibiting detectable expression under rapamycin (Rapa) treatment (mean ± s.d.). *n* = 25, 18, 16, 16, 20 tapes/cells for S1, S2, S3, S6, S7 respectively. Throughout this figure: the selected split site, S7, is highlighted in red.

**Figure S5.**
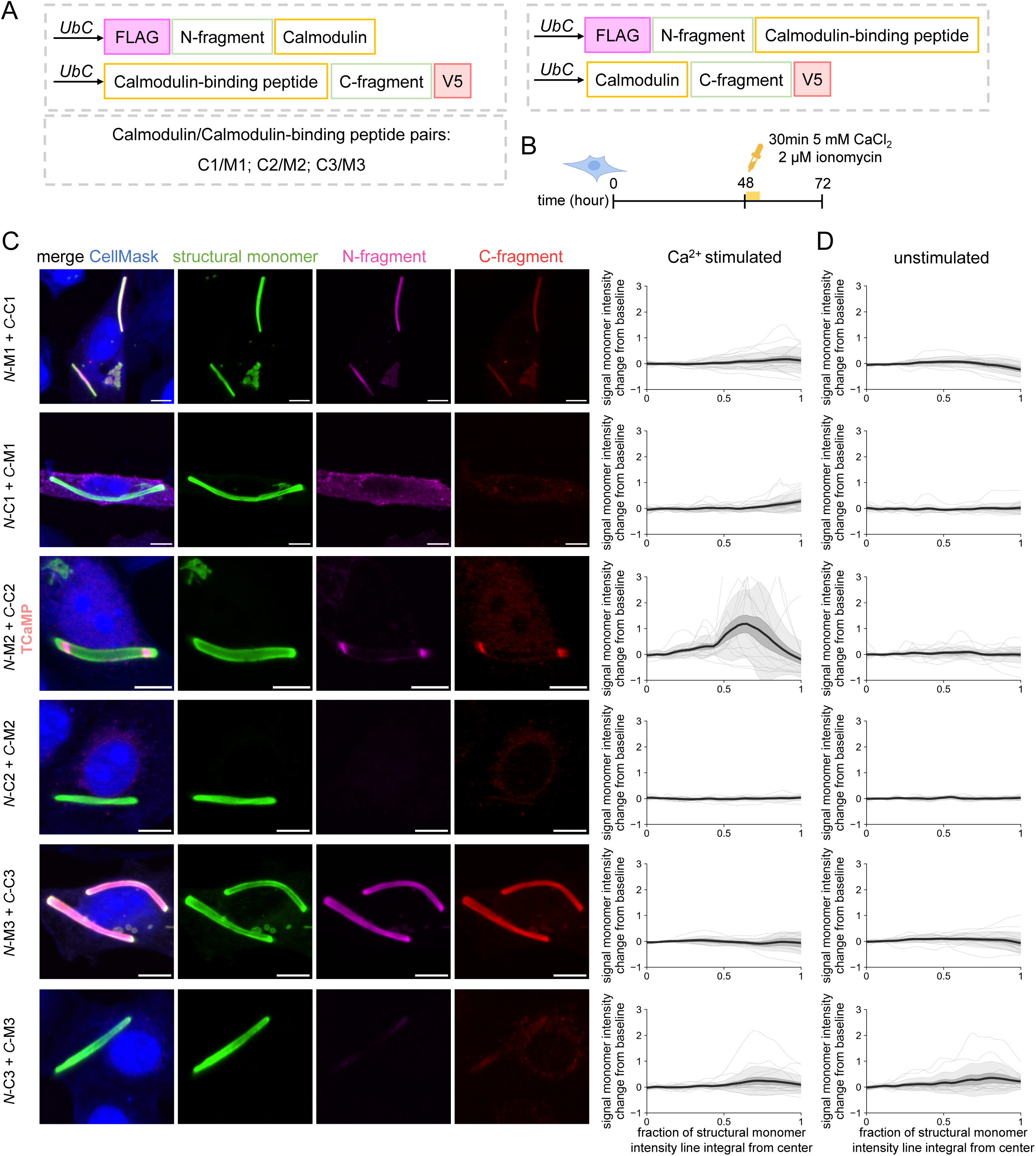
Screening CaM–CaM-binding peptide variants and their reciprocal configurations resulted in the TCaMP design. (**A**) Schematic of reciprocal split-monomer designs for engineering a Ca^2+^-dependent recorder. Calmodulin (CaM) and CaM-binding peptides were fused to either the N-or C-terminal fragments of the split monomer, generating two reciprocal orientations for each of three CaM/CaM-binding peptide pairs (C1/M1, C2/M2, and C3/M3). Linkers not shown. (**B**) Experimental timeline for screening Ca^2+^-dependent split-monomer designs in U2OS cells. Cells were transfected at 0 h, stimulated with 2 μM ionomycin and 5 mM CaCl_2_ for 30 min at 48 h, and fixed at 72 h. (**C**-**D**) Representative confocal images of tapes expressing the six reciprocal split-monomer designs following Ca^2+^ stimulation (**C**, left). Fluorescence profiles show signal monomer intensity changes from baseline plotted against the fraction of the structural monomer intensity line integral from the tape center for Ca^2+^ stimulated (**C**, right) and unstimulated control (**D**) groups. Center line, mean; dark shaded boundary, s.e.m.; light shaded boundary, s.d.; gray thin lines, data from individual tapes. The numbers of tapes (cells) in the unstimulated/Ca^2+^-stimulated groups were *n* = 11/15, 14/17, 16/15, 12/11, 15/7, and 11/16 for N-M1/C-C1, N-C1/C-M1, N-M2/C-C2, N-C2/C-M2, N-M3/C-C3, and N-C3/C-M3, respectively. Scale bars, 10 μm.

**Figure S6.**
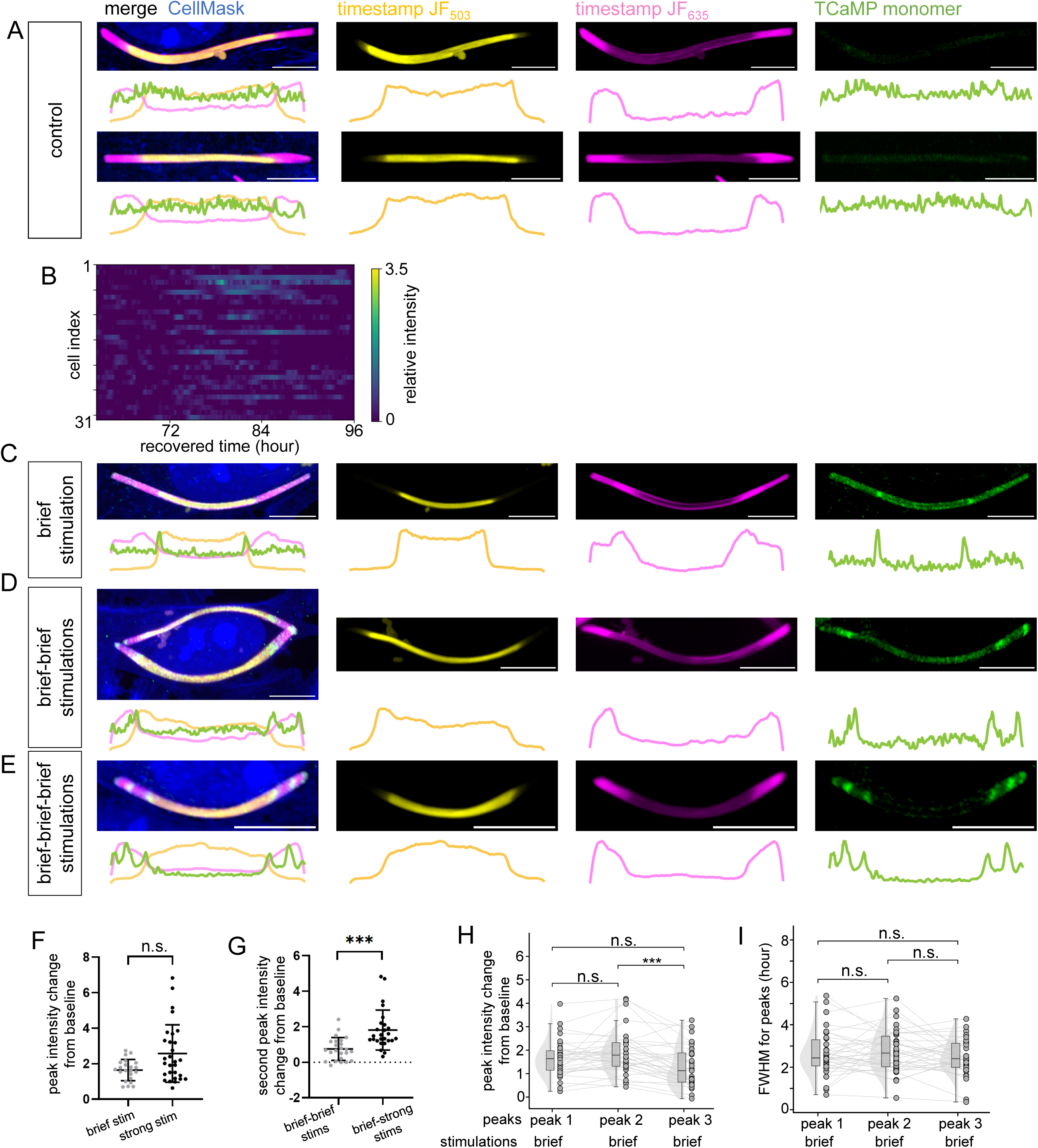
Representative TCaMP recordings under unstimulated and diverse Ca^2+^ stimulation conditions. (**A**) Representative confocal images and corresponding fluorescence intensity profiles of TCaMP tapes in unstimulated U2OS cells (control), with HaloTag-based timestamps defined as in **Figure 3E**. (**B**) Heatmap of individual TCaMP fluorescence intensity profiles plotted over recovered time for the unstimulated control group in **A**, with each row representing an individual tape (*n* = 31 tapes from 31 cells). (**C**–**E**) Representative confocal images and corresponding fluorescence intensity profiles of TCaMP tapes from the single brief Ca^2+^ stimulation (**C**), double brief–brief stimulation (**D**), and triple brief–brief–brief stimulation (**E**) groups, following the experimental paradigms described in **Figure 3E** and corresponding to the analyses in **Figure 3G**–**K**. (**F**) Quantification of peak TCaMP fluorescence intensity change from baseline following single brief or strong Ca^2+^ stimulation (mean ± s.d.). *n* = 22 and 29 tapes from 22 and 29 cells for brief and strong stimulations, respectively. (**G**) Quantification of the second peak fluorescence intensity change from baseline following sequential brief–brief or brief–strong Ca^2+^ stimulation (mean ± s.d.). *n* = 25 and 26 tapes from 25 and 26 cells for brief–brief and brief–strong stimulation, respectively. (**H**-**I**) Quantification of peak fluorescence intensity change from baseline (**H**) and full width at half maximum (FWHM; **I**) for three sequential TCaMP responses under brief-brief-brief triple stimulations, *n* = 34 tapes from 34 cells. Raincloud plots show data distributions, with half-violins representing density estimates, boxes indicating medians and IQRs, and whiskers extending to the most extreme values within 1.5 × IQR. Gray dots represent individual tapes, with thin gray lines connecting measurements from the same tape across the three peaks. Statistical significance was assessed using paired Wilcoxon signed-rank tests with Holm correction for multiple comparisons. Throughout this figure: n.s., not significant; **P < 0.01; ***P < 0.001; two-sided Mann–Whitney U test unless specified. Scale bars, 4 μm.

**Table S1.**
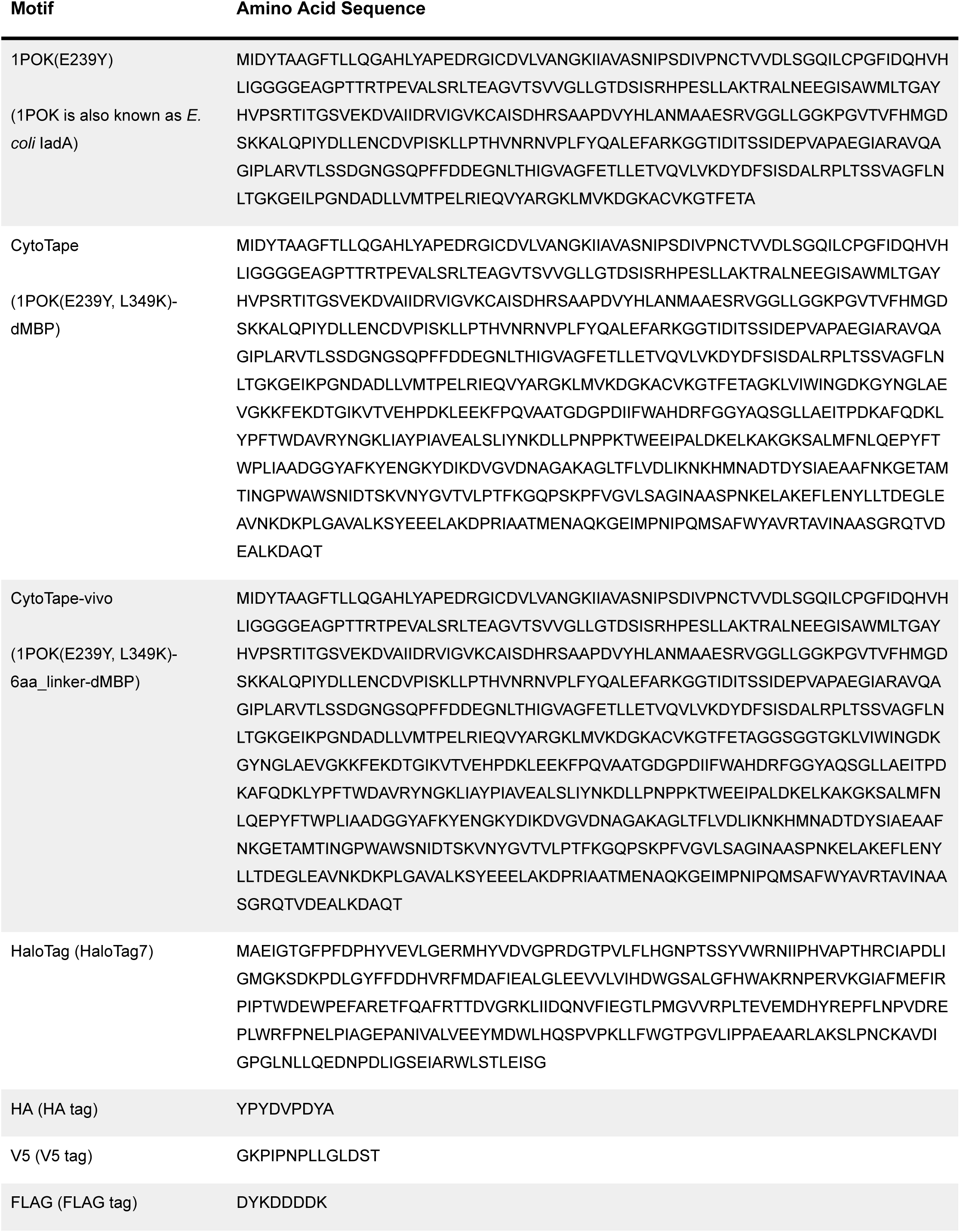
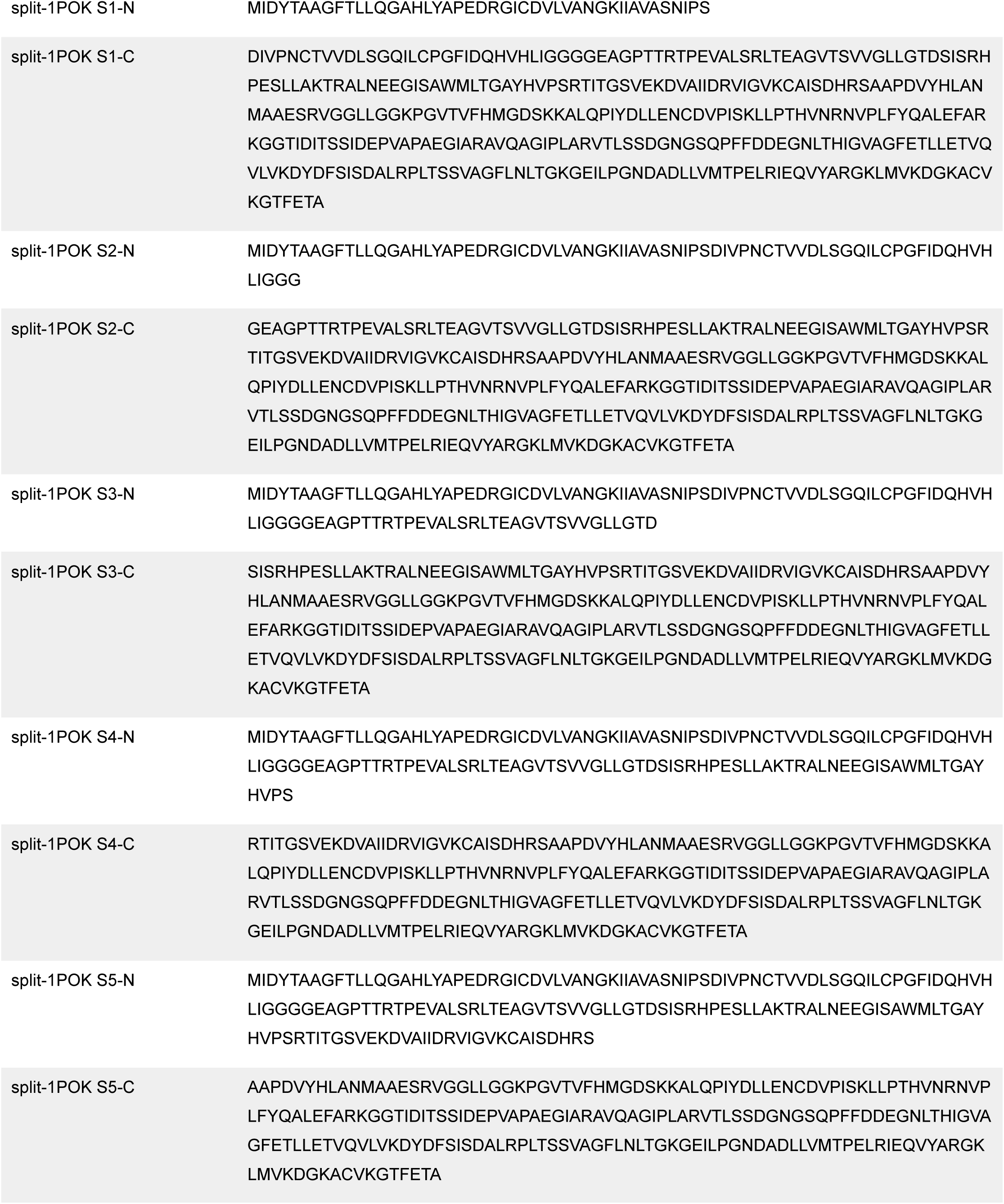
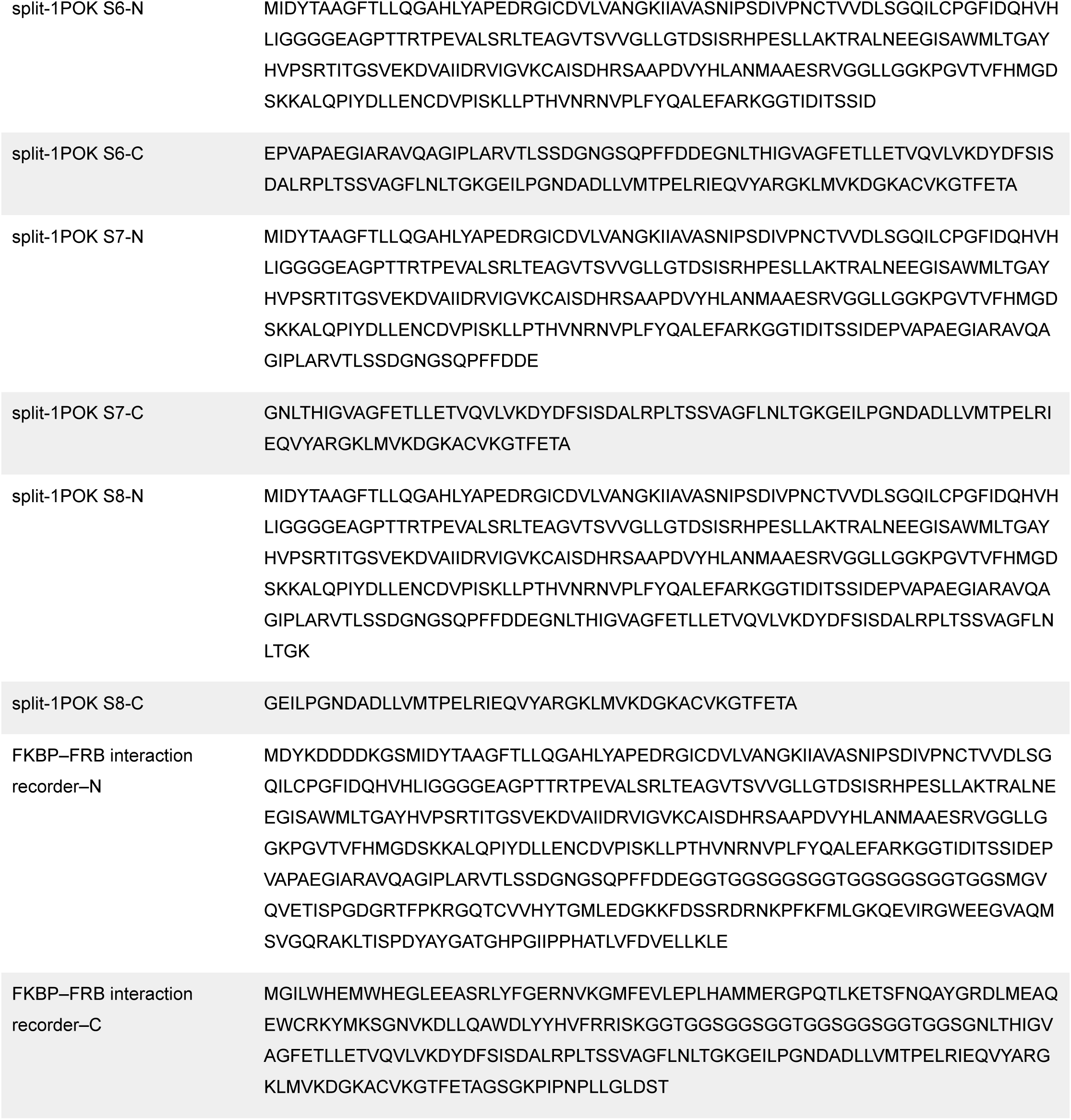
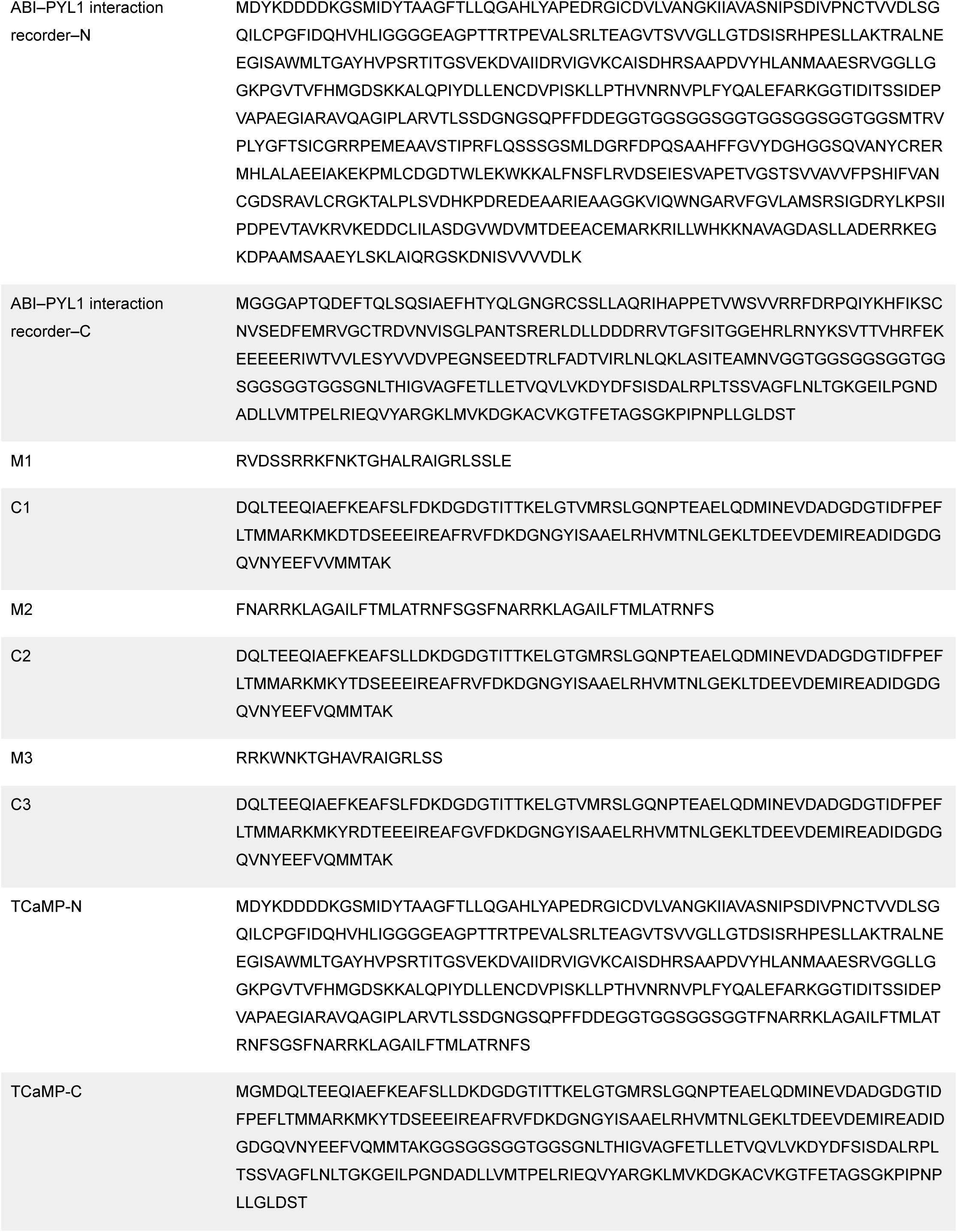
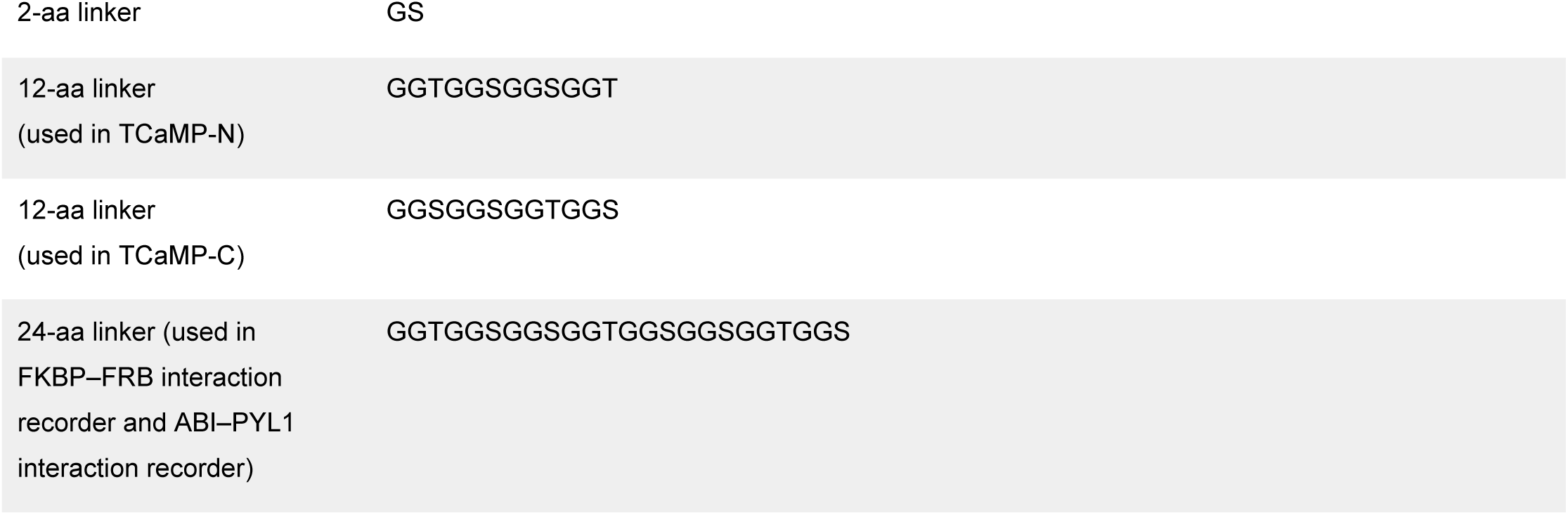
Sequences of genes and motifs used in this study.

## References

1. Yan, Y. et al. Genetically encoded systems for cytosolic activity recording and molecular archiving across time. 2026.06.15.732182 Preprint at 10.64898/2026.06.15.732182 (2026).

2. Clapham, D. E. Calcium Signaling. Cell 131, 1047–1058 (2007).

3. Nakai, J., Ohkura, M. & Imoto, K. A high signal-to-noise Ca2+ probe composed of a single green fluorescent protein. Nat. Biotechnol. 19, 137–141 (2001).

4. Chen, T.-W. et al. Ultrasensitive fluorescent proteins for imaging neuronal activity. Nature 499, 295–300 (2013).

5. Zhang, Y. et al. Fast and sensitive GCaMP calcium indicators for imaging neural populations. Nature 615, 884–891 (2023).

6. Fosque, B. F. et al. Labeling of active neural circuits in vivo with designed calcium integrators. Science 347, 755–760 (2015).

7. Wang, W. et al. A light- and calcium-gated transcription factor for imaging and manipulating activated neurons. Nat. Biotechnol. 35, 864–871 (2017).

8. Huppertz, M.-C. et al. Recording physiological history of cells with chemical labeling. Science 383, 890–897 (2024).

9. Linghu, C. et al. Recording of cellular physiological histories along optically readable self-assembling protein chains. Nat. Biotechnol. 41, 640–651 (2023).

10. Choi, J., Chen, J., Schreiber, S. L. & Clardy, J. Structure of the FKBP12-Rapamycin Complex Interacting with Binding Domain of Human FRAP. Science 273, 239–242 (1996).

11. Zheng, L. et al. Scalable and multiplexed recorders of gene regulation dynamics across weeks. Nature 652, 1038–1048 (2026).

12. Hosoi, H. et al. Rapamycin causes poorly reversible inhibition of mTOR and induces p53-independent apoptosis in human rhabdomyosarcoma cells. Cancer Res. 59, 886–894 (1999).

13. Liang, F.-S., Ho, W. Q. & Crabtree, G. R. Engineering the ABA Plant Stress Pathway for Regulation of Induced Proximity. Sci. Signal. 4, rs2–rs2 (2011).

14. Hyun, J. H. et al. Tagging active neurons by soma-targeted Cal-Light. Nat. Commun. 13, 7692 (2022).

15. Raffaello, A., Mammucari, C., Gherardi, G. & Rizzuto, R. Calcium at the Center of Cell Signaling: Interplay between Endoplasmic Reticulum, Mitochondria, and Lysosomes. Trends Biochem. Sci. 41, 1035–1049 (2016).

16. Ma, Q., Ye, L., Liu, H., Shi, Y. & Zhou, N. An overview of Ca^2+^ mobilization assays in GPCR drug discovery. Expert Opin. Drug Discov. 12, 511–523 (2017).

17. Brands, J. et al. A molecular mechanism to diversify Ca2+ signaling downstream of Gs protein-coupled receptors. Nat. Commun. 15, 7684 (2024).

18. Bird, G. S., DeHaven, W. I., Smyth, J. T. & Putney, J. W. Methods for studying store-operated calcium entry. Methods 46, 204–212 (2008).

19. de, M. F., et al. Human Golgi Antiapoptotic Protein Modulates Intracellular Calcium Fluxes. Mol. Biol. Cell 20, 3638–3645 (2009).

20. Lytton, J., Westlin, M. & Hanley, M. R. Thapsigargin inhibits the sarcoplasmic or endoplasmic reticulum Ca-ATPase family of calcium pumps. J. Biol. Chem. 266, 17067–17071 (1991).

21. Bootman, M. D., Berridge, M. J. & Taylor, C. W. All-or-nothing Ca2+ mobilization from the intracellular stores of single histamine-stimulated HeLa cells. J. Physiol. 450, 163–178 (1992).

22. Sagara, Y. & Inesi, G. Inhibition of the sarcoplasmic reticulum Ca2+ transport ATPase by thapsigargin at subnanomolar concentrations. J. Biol. Chem. 266, 13503–13506 (1991).

23. Yang, W. et al. The past, present and future of de novo protein design. Nature 652, 1139–1152 (2026).

24. Klewer, L. & Wu, Y.-W. Light-Induced Dimerization Approaches to Control Cellular Processes. Chem. – Eur. J. 25, 12452–12463 (2019).

25. Stanton, B. Z., Chory, E. J. & Crabtree, G. R. Chemically induced proximity in biology and medicine. Science 359, eaao5902 (2018).

26. Jozic, D., Kaiser, J. T., Huber, R., Bode, W. & Maskos, K. X-ray Structure of Isoaspartyl Dipeptidase from *E. coli*: A Dinuclear Zinc Peptidase Evolved from Amidohydrolases. J. Mol. Biol. 332, 243–256 (2003).

27. Abramson, J. et al. Accurate structure prediction of biomolecular interactions with AlphaFold 3. Nature 630, 493–500 (2024).

28. Pettersen, E. F. et al. UCSF ChimeraX: Structure visualization for researchers, educators, and developers. Protein Sci. 30, 70–82 (2021).

